# Chromatin-release of the long ncRNA A-ROD is required for transcriptional activation of its target gene DKK1

**DOI:** 10.1101/224360

**Authors:** Evgenia Ntini, Julia Liz, Jose M Muino, Annalisa Marsico, Ulf Andersson Ørom

## Abstract

Long non-coding RNAs (ncRNAs) are involved in both positive and negative regulation of transcription. Long ncRNAs are often enriched in the nucleus and at chromatin but whether chromatin-release plays a functional role is unknown. Here, we used epigenetic marks, expression level and strength of chromatin interactions to group long ncRNAs and find that those engaged in strong chromatin interactions are less enriched at chromatin in MCF-7 cells, suggesting a functional involvement of chromatin-release of long ncRNAs in transcriptional regulation. To study this further, we identify the long ncRNA *A-ROD*, an <u>a</u>ctivating <u>r</u>egulator <u>o</u>f the *Wnt* signaling inhibitor *<u>D</u>KK1.* We show that *A-ROD* enhances transcription elongation of *DKK1* in an RNA-dependent manner and that A-ROD recruits *EBP1* to the *DKK1* promoter. Our data suggest that the activating function depends on the release of A-ROD from chromatin, and further identify a functional regulatory interaction mediated by *A-ROD* in the transcription activation of *DKK1.* We propose that the release of a subset of long ncRNAs is important for their function, adding a new mechanistic perspective to the subcellular localization of long ncRNAs.

## Introduction

Long ncRNAs are functional RNAs not encoding proteins; they are distinct from short ncRNA species in that they are processed through the same molecular machinery as mRNAs and resemble mRNAs in their organization with several exons and introns as well as a 5' cap and frequent polyadenylation (Quinn & Chang, 2016). Functionally, long ncRNAs have been shown to bind proteins and direct their interaction with DNA (Vance et al, 2014; Wang et al, 2011b) or enhance their enzymatic activity (Lai et al, 2013; Marchese et al, 2016). In regulation of transcription, long ncRNAs are emerging as important factors with both positive and negative activity (Quinn & Chang, 2016). A subset of long ncRNAs has been shown to be expressed from enhancers and mediate the activation of transcription of adjacent target genes (Gomez et al, 2013; Lai et al, 2013; Li et al, 2016; Orom et al, 2010; Orom & Shiekhattar, 2013; Schaukowitch et al, 2014; Trimarchi et al, 2014; Wang et al, 2011b). Long ncRNAs are often enriched in the nucleus and in some cases tethered to chromatin suggesting an involvement in epigenetic regulation (Derrien et al, 2012; Khalil et al, 2009). While the nuclear localization of most long ncRNAs seems intuitive for their role in transcription regulation, it is not clear to which extent their activity depends on chromatin-association.

Enhancers are distant regulatory elements involved in regulation of target gene expression in a temporal- and/or cell type-specific manner. Enhancers can be identified using several epigenetic marks, including histone marks and DNA methylation. Functional enhancers have been described as defined transcription units with relatively elevated levels of the histone 3 lysine 4 trimethylation (H3K4me3) histone mark (Core et al, 2014; Pekowska et al, 2011) giving rise to long, spliced and polyadenylated ncRNAs which in turn mediate the enhancer function (Gomez et al, 2013; Lai et al, 2013; Orom et al, 2010; Trimarchi et al, 2014; Wang et al, 2011b). Enhancers can also be pervasively transcribed into relatively short and unstable enhancer-associated ncRNAs called eRNAs (Andersson et al, 2014; Kim et al, 2010; Wang et al, 2011a). The two groups, activating long ncRNAs and eRNAs, are distinct in terms of production, nature and stability, although a complete understanding of the functional repertoire of ncRNAs transcribed from active enhancers is not yet fulfilled (Orom & Shiekhattar, 2013; Quinn & Chang, 2016; Vucicevic et al, 2015).

Here, we address the chromatin-association of long ncRNAs by grouping them according to a combination of epigenetic marks and chromatin interactions. We find that actively expressed long ncRNAs, transcribed from loci engaged in RNA polymerase II (Pol II)-dependent chromatin interactions to active promoters, are less enriched in the chromatin fraction compared to other long ncRNAs expressed in human MCF-7 breast cancer cells. Using the long ncRNA A-ROD (Activating Regulator of DKK1) as an example, we show that transcription activation of its target gene DKK1 is accompanied by an A-ROD-dependent recruitment of the transcription factor EBP1 to the DKK1 promoter. The regulatory effect is exerted by A-ROD at its release from chromatin, suggesting that chromatin-release of activating long ncRNAs is necessary to mediate RNA-dependent regulation of their target gene expression.

## Results

### Expression and chromatin-release of long ncRNAs

Chromatin interactions are important for proper regulation of gene expression (Bulger & Groudine, 2011; Li et al, 2012). To determine the characteristics of long ncRNAs engaged in long-range chromatin interactions we used Pol II-dependent ChIA-PET data (Chromatin Interaction Analysis by Paired-End Tag sequencing) from MCF-7 cells (ENCODE, 2012; Li et al, 2012). ChIA-PET employs a chromatin immunoprecipitation step followed by high-throughput sequencing to detect long-range interactions at regions bound by a target protein of interest, in this case Pol II. Enriching for loci bound by a specific protein increases the probability to detect regulatory interactions and this technique has been used to map enhancer-promoter interactions (Heidari et al, 2014). To define our long ncRNA working dataset, we combined the ENCODE annotation of long ncRNAs, long ncRNAs reported in (Cabili et al, 2011) and *de novo* transcript assembly in MCF-7 cells from chromatin-associated RNA-sequencing data (Materials and Methods; GEO accession number GSE69507). To facilitate the analysis of chromatin marks at specific gene loci without confounding signals due to overlapping transcripts and to obtain high-confidence long ncRNAs we required transcripts to have at least one splicing junction, no coding potential (using CPAT) (Wang et al, 2013) and no overlap with annotated protein-coding genes. We obtain a list of 12,553 long ncRNAs transcribed from 10,606 genomic loci. This set was further narrowed down to 4,467 long ncRNAs with detectable expression in MCF-7 cells for the subsequent analyses (Supplementary Table S1, Figure 1A and Materials and Methods).

**Figure 1.**
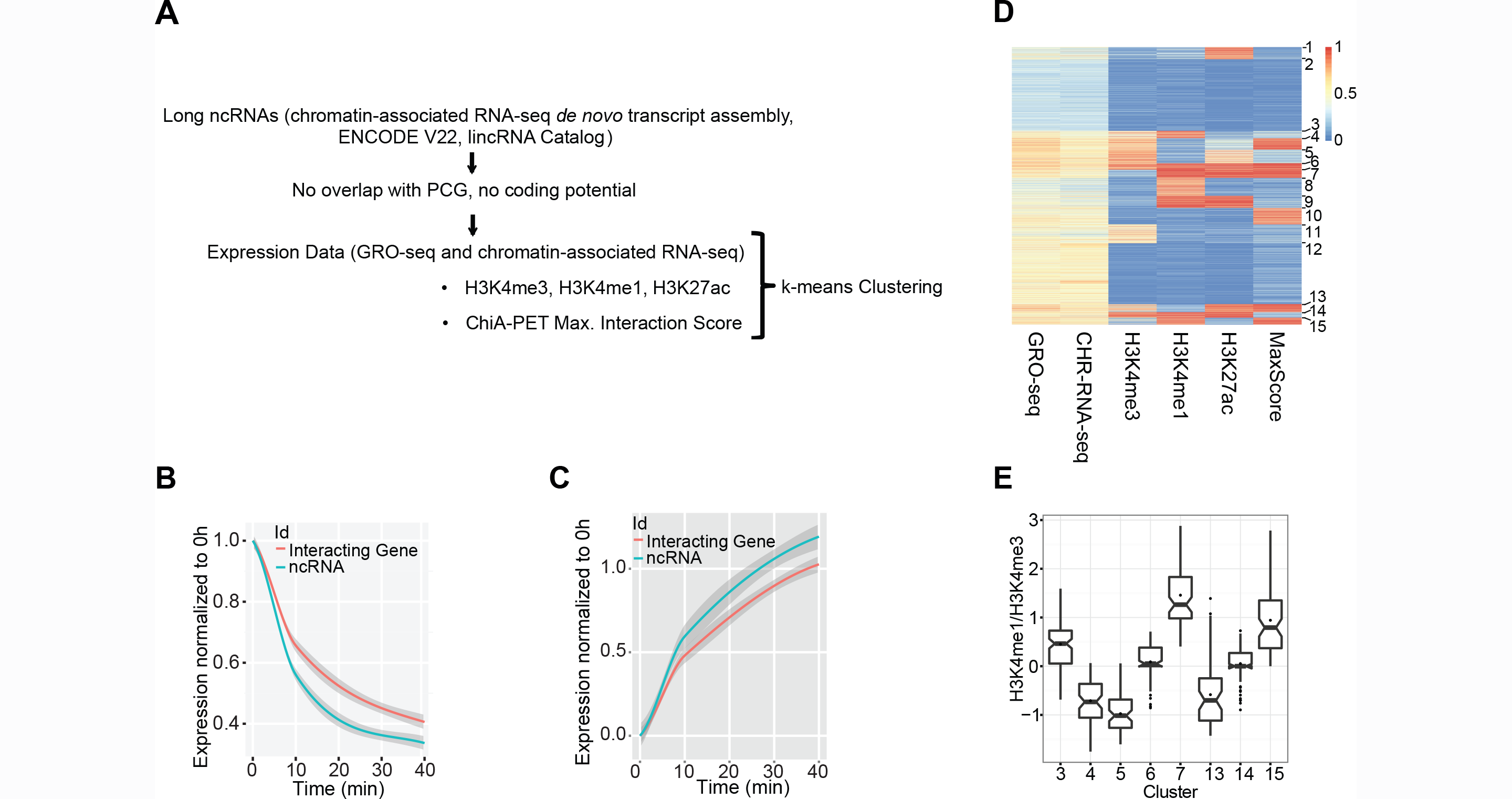
Grouping of long ncRNAs based on expression and epigenetic marks. (A) Schematic representation of the pipeline followed to define the final set of long ncRNAs used in this study and the k-means clustering parameters. (B) Estradiol mediated response for downregulated long ncRNA-interacting gene pairs (ChIA-PET interaction score > 200). GRO-seq RPKM from 10 and 40 min of E2 treatment are normalized to 0 h (EtOH control). Fitted regression model lines (loess curves) are drawn. The 95% confidence region is grey shaded. (C) Same as in B, for the estradiol-upregulated long ncRNA-gene pairs. *y*-axis is in the *log* scale. (D) K-means clustering of 4,467 long ncRNAs. Clustering parameters are histone mark signal values (ChIP-seq peaks from H3K4me3, H3K4me1 and H3K27ac in MCF-7), expression (GRO-seq and chromatin-associated RNA-seq log(RPKM)), and ChIA-PET maximum interaction scores. All values have been rescaled in the range 0 to 1 using min-max normalization. (E) H3K4me1/H3K4me3 ratio for relevant clusters, *y*-axis is in *log* scale.

The MCF-7 cell line is an estrogen receptor alpha (ERa) positive breast cancer cell line. Hormone treatment with estradiol (E2) induces broad transcriptional changes due to activation of the ERa. To examine the involvement of long ncRNAs in transcriptional regulation of target genes predicted by ChIA-PET under differential conditions we used expression data from control and 40 min E2 treated MCF-7 cells (GRO-seq) (Hah et al, 2013). We observe that the transcriptional response of long ncRNAs to E2 generally precedes the transcriptional response of the interacting genes (Figure 1B-C, EV1A). This is in agreement with previous studies reporting that transcriptional responses of stimuli-activated enhancers temporally precede expression changes of target genes (Arner et al, 2015; De Santa et al, 2010; Hah et al, 2011; Hsieh et al, 2014; Kim et al, 2015). Especially for the E2-downregulated long ncRNA/target gene pairs there is a lag in the time response of the long ncRNAs compared to target genes (Figure 1B), absent from random downregulated gene-gene pairs (Supplementary Figure S1A). This is in agreement with long ncRNAs being involved in positive regulation of their interacting target genes. In the case of E2-upregulated interacting pairs we see a similar but smaller lag (Figure 1C). This difference could be due to upregulated genes being significantly enriched in ERa-binding sites, which correlates with an increase in expression (Supplementary Figure S1B), triggered as a direct response to ERa binding.

We grouped the long ncRNAs using epigenetic marks associated with transcription regulation by incorporating H3K4 monomethylation (H3K4me1) (GEO accession number GSM588569); H3K27 acetylation (H3K27Ac) and H3K4me3 (ENCODE, 2012); RNA expression (GRO-seq, GEO accession number GSE43835) (Hah et al, 2013) and chromatin-associated RNA-seq, DNA methylation at long ncRNA promoters, as well as Pol II ChIA-PET interaction data from MCF-7 cells (ENCODE, 2012; Li et al, 2012). Using ENCODE DNA methylation data we find a correlation between promoter DNA hypomethylation and long ncRNA expression specificity across nine cell lines, suggesting an involvement of DNA methylation status in long ncRNA expression (Supplementary Figure S1C). We extended the dataset of long ncRNA promoters with assayed DNA methylation in MCF-7 cells by array-based DNA bisulfite sequencing (GEO accession number GSE69507) (Supplementary Figure S1D). We then performed k-means clustering of 2,685 long ncRNA transcripts according to epigenetic marks and transcription, where all data are available, to group the long ncRNAs into 15 clusters (Supplementary Figure S2A). Importantly, highly expressed long ncRNAs associating with enhancer marks show low DNA methylation and strong ChIA-PET interactions (Supplementary Figure S2A). We therefore reduced the parameters by excluding DNA methylation to extend the k-means clustering analysis to all 4,467 long ncRNAs of our initial dataset (Figure 1D). Clusters with strong ChIA-PET interactions are marked with promoter (high H3K4me3) or enhancer (high H3K4me1 and/or H3K27ac) histone marks. Of particular interest for our study of activating long ncRNAs, cluster 6 contains 104 long ncRNAs showing enrichment of all included histone marks (Figure 1D and Supplementary Figure S2B) and relatively high expression. As expected, the promoters of the long ncRNAs of cluster 6 with assayed DNA methylation are commonly hypomethylated (Supplementary Figure S2B). The cluster shows strong chromosomal interactions between long ncRNAs and target genes and the E2-mediated transcription changes of the long ncRNAs and ChIA-PET predicted target genes correlate well (Supplementary Figure S2C). In addition, the intermediate H3K4me1/H3K4me3 ratio of cluster 6 is characteristic of actively transcribed enhancers (Figure 1E) (Vucicevic et al, 2015). The positional distribution of histone marks shows characteristic bidirectional deposition around transcription start sites (TSS), with cluster 6 showing on average the highest levels of H3K27Ac, a hallmark of active enhancers (Supplementary Figure S2D–F).

Long ncRNAs are enriched in chromatin compared to mRNA (Derrien et al, 2012; Werner & Ruthenburg, 2015) and this is thought to reflect their nuclear function. We calculated the chromatin-association of long ncRNAs in MCF-7 cells by normalizing chromatin-associated RNA-seq to ENCODE available nuclear polyA+ RNA-seq. Interestingly, long ncRNAs of clusters with higher ChIA-PET scores (median = 1 and mean > 0.9) (Clusters 4, 6, 7, 13 and 15) are significantly less enriched in the chromatin fraction (Wilcoxon-Mann-Whitney test p-value < 2.2e-16) (Figure 2A-B), suggesting that chromatin-dissociation could play a functional role for long ncRNAs transcribed from loci engaged in strong chromatin interactions. To exclude any bias in the analysis by using the nuclear polyA+ data for normalization, we sequenced total nucleoplasmic RNA (after depletion of ribosomal species) and re-assessed chromatin-association as the ratio of total chromatin-associated to nucleoplasmic RNA. This results in an overall similar distribution, with clusters 4, 6 and 13 of strong chromatin interactions showing on average lower chromatin association (Supplementary Figure S3A). Because of the observed difference in the distribution by using nuclear polyA+ data (Figure 2B) we conducted further analysis to investigate what defines the chromatin-association of long ncRNAs. We performed linear regression with the parameters GRO-seq, H3K4me3, H3K4me1, ChIA-PET and H3K27Ac (Supplementary Figure S3B). We see that the ChIA-PET interaction strength and the deposition of H3K4me3 produce the highest negative coefficients (−2.559 and −2.856, respectively, at p-value < 2.2e-16) in predicting chromatin-association with a fairly good linear fit (correlation = 0.476). The negative sign of the coefficients means a significant negative contribution, i.e. strong ChIA-PET interactions and H3K4me3 result in low chromatin association of long ncRNAs and thus a relatively higher release. By including nuclear polyA+ expression as an extra parameter in the linear regression fit, the correlation improves only slightly (correlation = 0.487), with ChIA-PET interaction strength and H3K4me3 remaining the most significant parameters in determining the fit (Supplementary Figure S3C). We also employed Principal Component Analysis (PCA) to compare chromatin-enriched (log2 chromatin-association > 0) versus nucleoplasmic-enriched (log2 chromatin-association < 0) long ncRNAs, using the same parameters as for the linear regression (Figure 2C). In PCA, the five variables are collapsed into a two-dimensional representation in the two Principal Components (PC) PC1 and PC2. Each PC occupies one dimension and the midpoint has value zero. The positive or negative sign of the variables defines the direction of a given variable on a single dimension PC vector. The bigger the (absolute) value of a variable, the bigger the impact it has (either positive or negative contribution) in explaining the variation for that PC. The PCA result shows that most of the nucleoplasmic enriched long ncRNAs have a PC1 score > 0 (right of the midpoint, blue dots) while grouping distinctly from chromatin-enriched long ncRNAs (PC1 score < 0, left of the midpoint, red dots). The parameters ChIA-PET interaction strength and H3K4me3 are the major contributors (Figure 2C). This is in agreement with the result of the linear regression analysis. We then performed differential expression analysis using DESeq2 (Love et al, 2014) to extract significantly chromatin-enriched versus nucleoplasmic-enriched long ncRNAs (at p-adjusted < 0.1; Supplementary Tables S2 and S3). Importantly we see that nucleoplasmic-enriched long ncRNAs are engaged in strong ChIA-PET interactions with significantly higher scores when compared to chromatin-enriched long ncRNAs or to long ncRNAs not enriched in either of the two fractions (Wilcoxon-Mann-Whitney p-value = 9.462e-16) (Figure 2D-E). Together these data support that long ncRNAs engaged in strong chromatin interactions show significantly lower chromatin-association, suggesting that chromatin-release is important for the function of these long ncRNAs.

**Figure 2.**
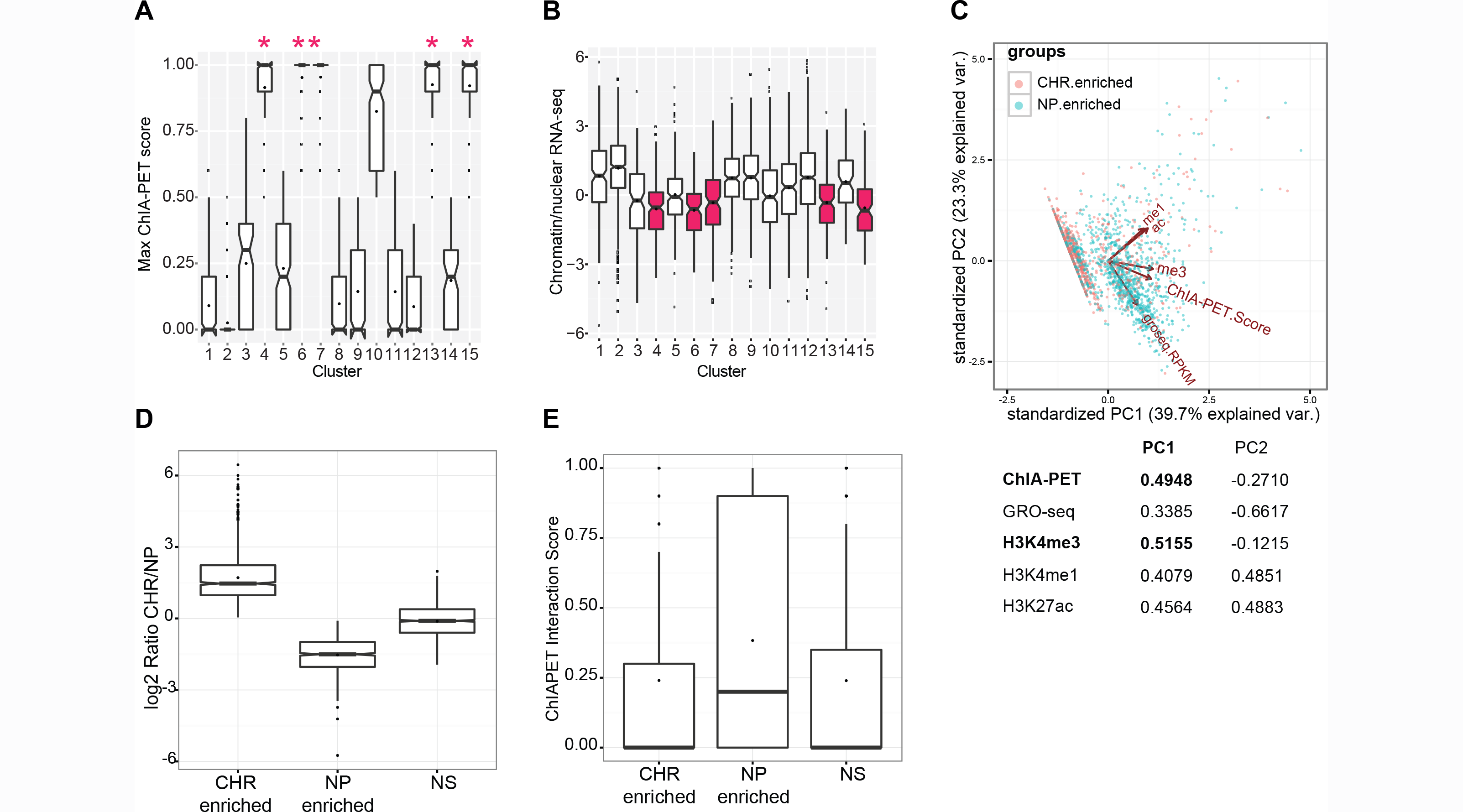
Chromatin-association of long ncRNAs. (A) Comparison of ChIA-PET interaction scores for all clusters. Marked are the clusters with the strongest interactions (score median at 1 and mean > 0.9). (B) Comparison of the long ncRNA chromatin-association character for all clusters. The ratio chromatin-associated RNA-seq RPKM to nuclear polyA+ RPKM is plotted in the *log* scale. (C) Principal component analysis (PCA) applied to chromatin-enriched (log2 chromatin-association > 0) and nucleoplasmic-enriched (log2 chromatin-association < 0) long ncRNAs. The used variables (ChIA-PET, GRO-seq, H3K4me3, H3K4me1, H3K27Ac) and their contribution in explaining the variance within each PC are included in the table underneath. (D) Boxplot showing the *DESeq2* derived log2 fold changes in expression using total chromatin-associated versus nucleoplasmic RNA-seq, for 1154 significantly chromatin-enriched long ncRNAs (at *DESeq2* p-adjusted < 0.1; Supplementary Table S2), 968 significantly nucleoplasmic-enriched long ncRNAs (Supplementary Table S3) and 1807 long ncRNAs not enriched in either of the two fractions (non-significant, ‘NS’). (E) Boxplot showing distribution of ChIA-PET interaction scores for the three long ncRNA sets defined as in (D). Nucleoplasmic-enriched long ncRNAs (‘NP enriched’; Supplementary Table S3) show on average significantly higher ChIA-PET interaction scores (Wilcoxon-Mann-Whitney test p-value = 9.462e-16 to ‘CHR enriched’ and p-value < 2.2e-16 to ‘NS’).

### A-ROD as a tissue-specific regulator of DKK1 expression

We further examined cluster 6 due to the prominent presence of enhancer marks and high chromatin-dissociation. We ranked the long ncRNA-interacting gene pairs of cluster 6 according to fold changes in expression upon 40 min E2-treatment versus control in MCF-7 cells. We focused on downregulated interacting pairs without ERa binding sites determined by ChIP-seq (Welboren et al, 2009) +/-20 kb from the TSS to avoid direct transcriptional responses to E2. The top candidate is the long ncRNA *A-ROD* transcribed 130 kb downstream of Dickkopf homolog 1 (DKK1) on the opposite strand (Supplementary Figure S4A). The DKK1 protein is involved in regulating the Wnt signaling pathway in many tissues and in breast cancer (Mikheev et al, 2008; Niida et al, 2004; Qiao et al, 2008). Downregulation of A-ROD precedes that of DKK1 upon E2-treatment (Figure 3A) in accordance with the general faster transcriptional response of long ncRNAs shown in Figure 1B. A-ROD is a multi-exonic long ncRNA that is polyadenylated and relatively highly expressed (Supplementary Figure S4A), suggesting that it is a bona fide long ncRNA transcribed from an active enhancer rather than an enhancer-associated eRNA (Lam et al, 2014). The expression of A-ROD in MCF-7 cells is high compared to other cell lines included in the analysis (Figure 3B) and generally shows an expression restricted to epi- and endothelial cells (NHEK and HMEC; ENCODE, 2012). Using data from the Genotype-Tissue Expression Project (GTEx), we observe a correlation between DKK1 and A-ROD expression among several tissue samples (Supplementary Figure S4B). In addition, there is a strong and significant quantitative difference in DKK1 expression between A-ROD–positive and A-ROD–negative samples (Wilcoxon-Mann-Whitney p-value < 2.2e-16; Figure 3C), as well as a significant correlation between DKK1 and A-ROD expression across a panel of both ERa-positive and ERa-negative breast cancer samples (data from the Cancer Genome Atlas) (Figure 3D). Taken together these data support A-ROD as a tissue-specific activator of DKK1 expression.

**Figure 3.**
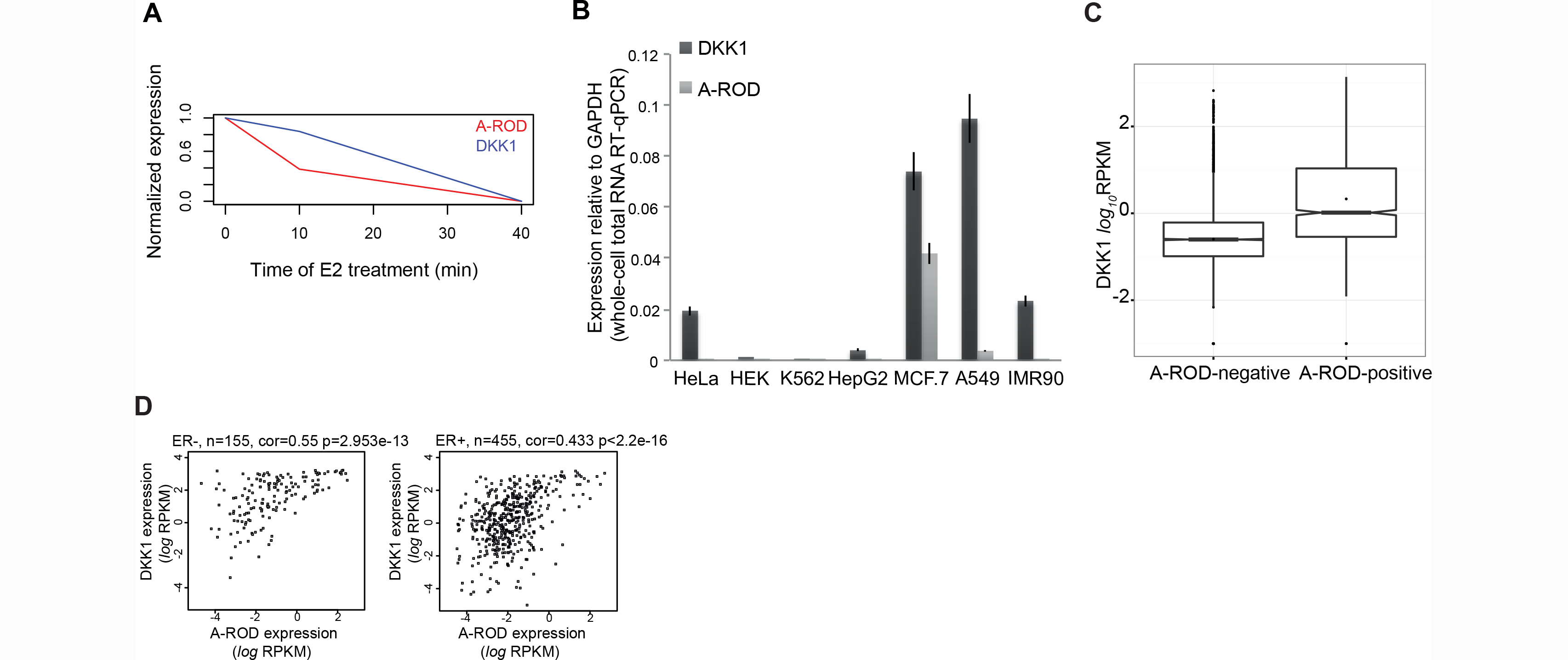
A-ROD is a tissue-specific regulator of DKK1 expression. (A) A-ROD and DKK1 expression as a time response to E2 induction. GRO-seq RPKM from 0 h (EtOH control), 10 and 40 min of E2 treatment were rescaled in the range 0 to 1 using min-max normalization. (B) Expression of A-ROD and DKK1 measured by RT-qPCR across seven cell lines, normalized to GAPDH. (C) Expression of DKK1 in A-ROD–positive (n = 1659) and A-ROD–negative tissue samples (n = 6896) using GTEx data. Wilcoxon-Mann-Whitney test p-value < 2.2e-16. (D) Correlation of A-ROD and DKK1 expression (total exon RPKM in the *log* scale) in ER positive and ER negative breast cancer samples. Data obtained from the Cancer Genome Atlas.

### A-ROD activates DKK1 transcription

Knock-down of enhancer-like long ncRNAs using siRNAs has been shown to affect the expression of their target genes (Lai et al, 2013; Orom et al, 2010; Trimarchi et al, 2014; Wang et al, 2011b). Upon knock-down of A-ROD using different siRNA sequences in MCF-7 cells, there is a reduction in DKK1 mRNA levels comparable to the extent of A-ROD depletion (Figure 4A). siRNA-mediated depletion of DKK1 does not affect A-ROD levels (Figure 4A). Neither A-ROD nor DKK1 siRNA-mediated depletion affect the transcript level of two independent long ncRNAs. To further determine whether A-ROD regulates DKK1 at the transcriptional level, we performed ChIP for Pol II in control and siRNA treated cells, using the N-20 antibody (Figure 4B). We see a significant decrease in total Pol II association at the DKK1 gene body, whereas there is no significant change in total Pol II occupancy at either GAPDH or A-ROD loci, supporting a specific regulatory effect of A-ROD on DKK1 transcription. These data are in agreement with a direct transcriptional effect on DKK1 mediated by the long ncRNA A-ROD.

**Figure 4.**
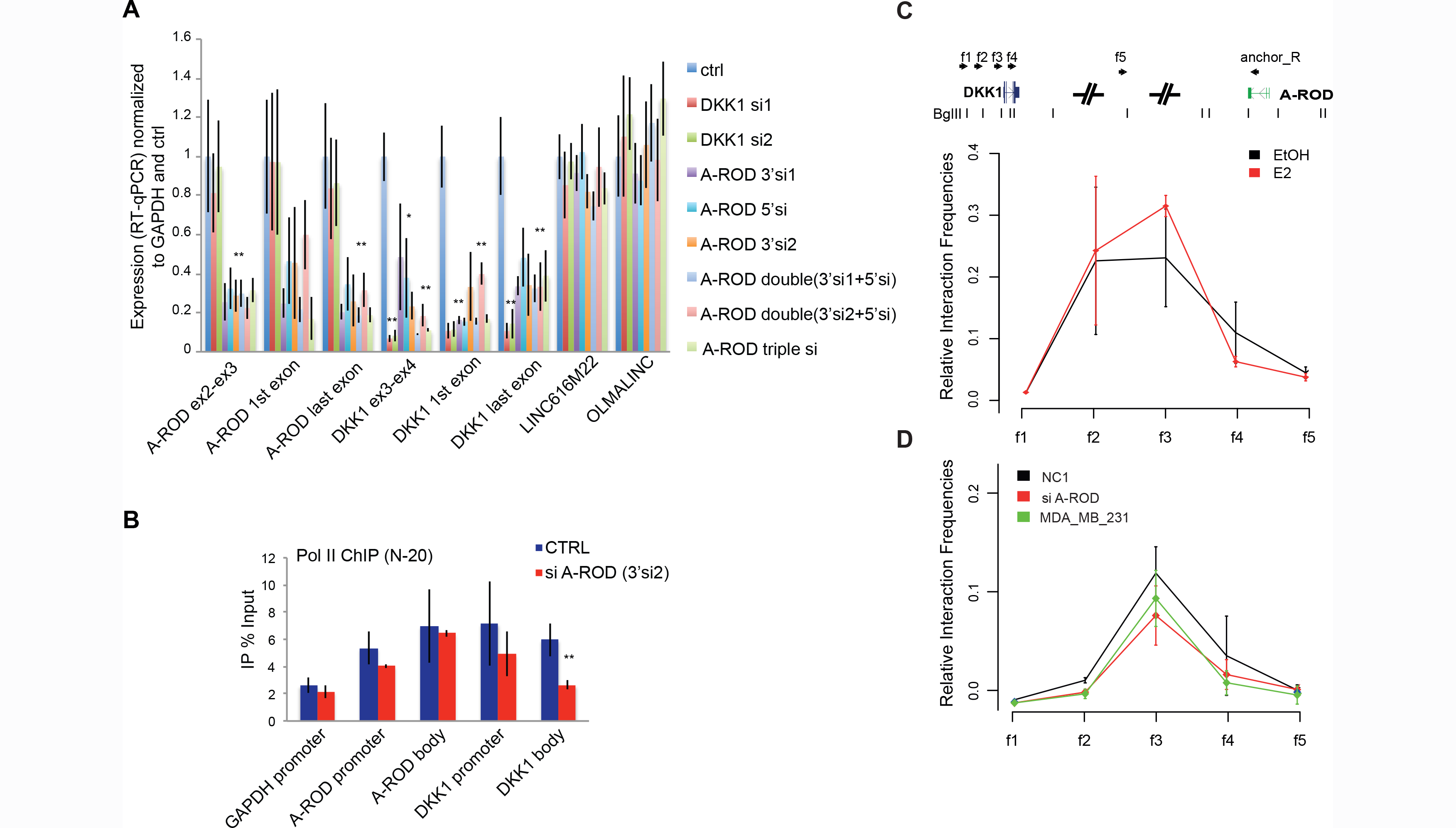
A-ROD enhances DKK1 transcription within the context of pre-established chromosomal interactions. (A) RT-qPCR levels of A-ROD, DKK1 and two independent long ncRNAs, in control and various siRNA treated cells, normalized to GAPDH. Error bars represent standard deviations from three independent experiments. (B) ChIP assaying total Pol II occupancy in control and A-ROD siRNA (#A-ROD.3’si2) treated MCF-7 cells using the N-20 antibody. Error bars represent standard deviations from three independent experiments. (C and D) Chromosome conformation capture of (C) the DKK1 A-ROD interaction in control (EtOH) and 40 min E2 induced MCF-7 cells and of (D) the DKK1 A-ROD interaction in control (NC1) or A-ROD siRNA transfected MCF-7 cells and in MDA-MB-231 cells. Relative interaction frequencies derived as described in Supplementary Methods.

### Long ncRNA target specificity is ensured within the physical proximity provided by pre-established chromosomal interactions

Enhancers are looped to promoters of regulated genes in the genome through long-range chromosomal interactions (Bulger & Groudine, 2011). The effect of long ncRNAs in determining the interaction landscape is not firmly established. Although long ncRNAs have been described to affect the long-range chromosomal interaction when associated with the subunit 12 of the Mediator complex (Lai et al, 2013), other studies report no effect on chromatin looping (Trimarchi et al, 2014; Wang et al, 2011b). This suggests that several mechanistic scenarios may exist for the large, diverse class of long ncRNAs. We performed chromosome conformation capture (3C) in control and E2-treated MCF-7 cells to assess the effects of A-ROD expression level and transcriptional changes on chromatin interactions to the DKK1 promoter. We also used another cell line, MDA-MB-231, where DKK1 but not A-ROD is expressed (Supplementary Figure S4A). The promoter of A-ROD in MDA-MB-231 cells is methylated (not shown) and does not show characteristics of active transcription as opposed to MCF-7 cells where the promoter is hypomethylated and A-ROD highly expressed. While we can recapitulate the chromatin interaction between A-ROD and DKK1 in MCF-7 cells by 3C, E2-treatment does not affect this interaction despite transcriptionally repressing both A-ROD and DKK1 (Figure 4C). Additionally, knock-down of A-ROD does not result in changes in the interaction despite significant effects on transcription of DKK1 (Figure 4D), supporting that the RNA does not play an active role in establishing or maintaining the chromatin interaction. In the MDA-MB-231 cell line we observe chromatin interactions between the DKK1 and A-ROD loci comparable to MCF-7 cells (Figure 4D) despite no active transcription of A-ROD. These data argue that neither the RNA nor the transcription affects this interaction, and that the chromatin interactions are established prior to the regulation mediated by A-ROD on DKK1.

### Regulation occurs independent of chromatin-bound A-ROD

While long ncRNAs associate to a higher degree with chromatin than mRNAs (Derrien et al, 2012; Werner & Ruthenburg, 2015), to which extent their activity depends on this association is not known. Analysis of chromatin-association of different groups of long ncRNAs based on epigenetic marks suggests that chromatin-release is involved in the mechanism of function of long ncRNAs engaged in strong chromatin interactions (Figure 2A-B). To address whether the regulatory effect of the long ncRNA happens while associated to chromatin or at or after dissociation from the chromatin-associated site of transcription, we fractionated MCF-7 cells, treated with either control or A-ROD siRNA, as described in Supplementary Methods (Conrad et al, 2014) (Supplementary Figure S5), and analyzed the nucleoplasmic and chromatin-associated RNA. While the chromatin-associated fraction is poorly targeted by siRNAs (Figure 5A), the nucleoplasmic A-ROD can be readily targeted by siRNA (Figure 5B), in agreement with recent studies showing assembly and function of the siRNA machinery in the nucleus (Gagnon et al, 2014). Interestingly, upon A-ROD siRNA transfection we can recapitulate the decrease in steady-state DKK1 mRNA levels we see from whole cells (Figure 4A) for the nucleoplasmic form of DKK1, but not for the chromatin-associated fraction (Figure 5A-B). This result could suggest that a repressive transcriptional effect caused by downregulation, for instance at the transcription elongation step, may not be captured as an efficient decrease in the steady-state chromatin-associated RNA levels, whereas the result becomes evident in the steady-state nucleoplasmic and cytoplasmic RNA fractions where effective turn-over takes place in the long term. To further unveil the mechanistic details of this regulation, and while considering that downregulation of transcription may not be properly reflected in steady-state chromatin-associated RNA, we purified nascent RNA, labeled by 5-bromouridine (BrU) incorporation from the chromatin fraction following cell fractionation in control and A-ROD siRNA transfected cells. We used this approach to study knock-down of nascent A-ROD RNA as well as transcription effects on DKK1. Analysis of nascent RNA obtained after a 15 minutes pulse shows a small effect of A-ROD siRNA on A-ROD levels as determined by qPCR of the 1st exon (Figure 5C). There is however a significant reduction when quantifying the last exon arguing that the fully transcribed A-ROD can be targeted by siRNAs while still loosely associated to the chromatin template, or at the very point of its release from chromatin. This result, recapitulated using an siRNA sequence targeting either the 3’ or the 5’ of the A-ROD transcript, supports that A-ROD RNA is responsible for the transcriptional activation of DKK1 while loosely associated to chromatin or at its release from the chromatin-associated site of transcription. We used these data to calculate the transcription pausing index (Adelman & Lis, 2012; Core et al, 2008; Hah et al, 2011). Upon A-ROD knock-down there is a significant increase in the pausing index of the DKK1 locus (Figure 5D), in agreement with A-ROD being involved in transcriptional enhancement of DKK1 expression.

**Figure 5.**
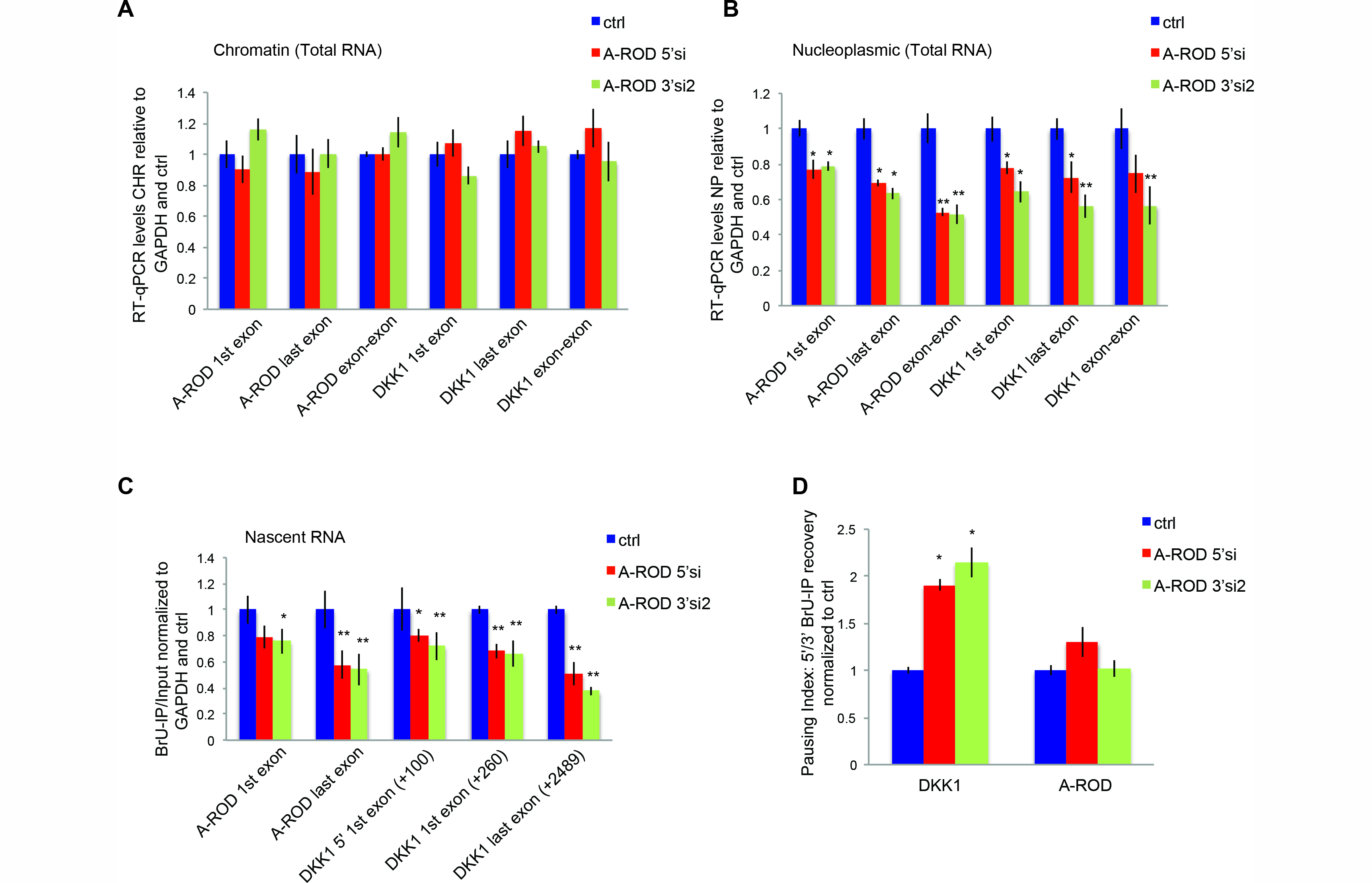
A-ROD exerts transcriptional regulatory activity at its release from chromatin. (A-B) Normalized RT-qPCR expression levels of A-ROD and DKK1 upon cell fractionation, in the chromatin-associated (A) and nucleoplasmic fraction (B), in control (ctrl) and siRNA-treated MCF-7 cells. Two different siRNA sequences targeting either the 5’ (A-ROD.5’si) or the 3’ (A-ROD.3’si2) of A-ROD were used. Error bars represent standard deviations from three independent experiments. (C) Expression levels of A-ROD and DKK1 in the BrU-labeled nascent RNA (BrU IP done from the chromatin-associated RNA fraction) measured by RT-qPCR in control and A-ROD siRNA treated cells, assayed by two different siRNA sequences targeting either the 5’ (A-ROD.5’si) or 3’ (A-ROD.3’si2) of A-ROD. Values were normalized to GAPDH exon and control. (D) Transcriptional pausing index for A-ROD and DKK1 assessed by extracting the ratio 5’/3’ IP recovery (BrU-IP over Input for two distinct 5’ and 3’ RT-qPCR amplicons) in control and A-ROD siRNA treated cells.

### A-ROD implication in regulation of DKK1 transcription elongation at its release from chromatin

Taken together, the above results from the siRNA-mediated depletion of A-ROD suggest that the chromatin-associated form of A-ROD is not responsible for DKK1 transcriptional enhancement. To further substantiate that it is rather the chromatin-released form of A-ROD exerting the regulatory effects, we used RNaseH-active antisense oligonucleotides (ASO’s) targeting intronic sequences of A-ROD for chromatin-associated knock-down. We assayed three different ASO sequences (provided in the Supplementary Materials and Methods) one of which showed efficient knock-down of chromatin-associated A-ROD (Figure 6A). Following cell fractionation at an early time-point (16h post-transfection), we assessed the chromatin-associated and nucleoplasmic levels of A-ROD by RT-qPCR. A-ROD levels appear significantly reduced in the chromatin fraction, assessed by measuring different amplicons, whereas the steady-state nucleoplasmic levels of A-ROD remain unaffected (Figure 6A-B). This is not surprising as the transfected ASO is targeting purely intronic sequence, and the analysis is done only 16h post-transfection. Yet ASO transfection causes an accompanying repressive transcriptional effect on DKK1 expression (Figure 6C-D) similar to the one we see upon siRNA transfection (Figure 5C-D), where the steady-state chromatin-associated levels of A-ROD remain unaltered (Figure 5A). These results fit together in a model where what is actually depleted by the intronic ASO is the active form of A-ROD at the point of its nascent transcript release from the chromatin-associated template. In a model where the intronic ASO interferes with and impedes the efficiency of co-transcriptional processing (splicing) of A-ROD, this would consequently result in reduced levels of the released (spliced) active form of A-ROD. To examine this we performed BrU-IP from the chromatin-associated and nucleoplasmic fraction in control and 16h ASO treated cells, and extracted the ratio of nascent nucleoplasmic to nascent chromatin-associated (or loosely associated to chromatin) A-ROD. The result supports that the intronic ASO affects the release of the nascent A-ROD transcript, whereas there is no effect on nascent GAPDH transcript release (Supplementary Figure S6A). The reduction of nascent DKK1 release is likely due to the consequent repressive effect on transcription caused by the depletion of the active form of A-ROD. This is in agreement with a scenario where active transcription links to efficient nascent transcript release. Depletion of the active form of A-ROD via ASO transfection causes downregulation of DKK1 transcription (Figure 6C), likely exerted at the transcription elongation step, as suggested by the increase in the pausing index (Figure 6D). To further substantiate an effect on transcription elongation we performed ChIP of Pol II phosphorylated at Ser2 (phospho-Ser2 Pol II), which reflects the transcriptionally engaged RNA Pol II (Adelman & Lis, 2012). There is a significant decrease of phospho-Ser2 Pol II along the DKK1 body (Figure 7A) and an accompanied increase in the pausing index (Figure 7B), when knocking-down A-ROD with either ASO or a pool of siRNAs. Furthermore, A-ROD depletion does not alter the TFIIB ChIP signal at the DKK1 promoter, suggesting that it does not affect the formation of pre-initiation complexes (PIC) at the DKK1 promoter (Supplementary Figure S6B). Taken together the above results support that A-ROD mediated transcriptional enhancement of DKK1 is exerted at the level of productive elongation rather than at the transcription initiation step.

**Figure 6.**
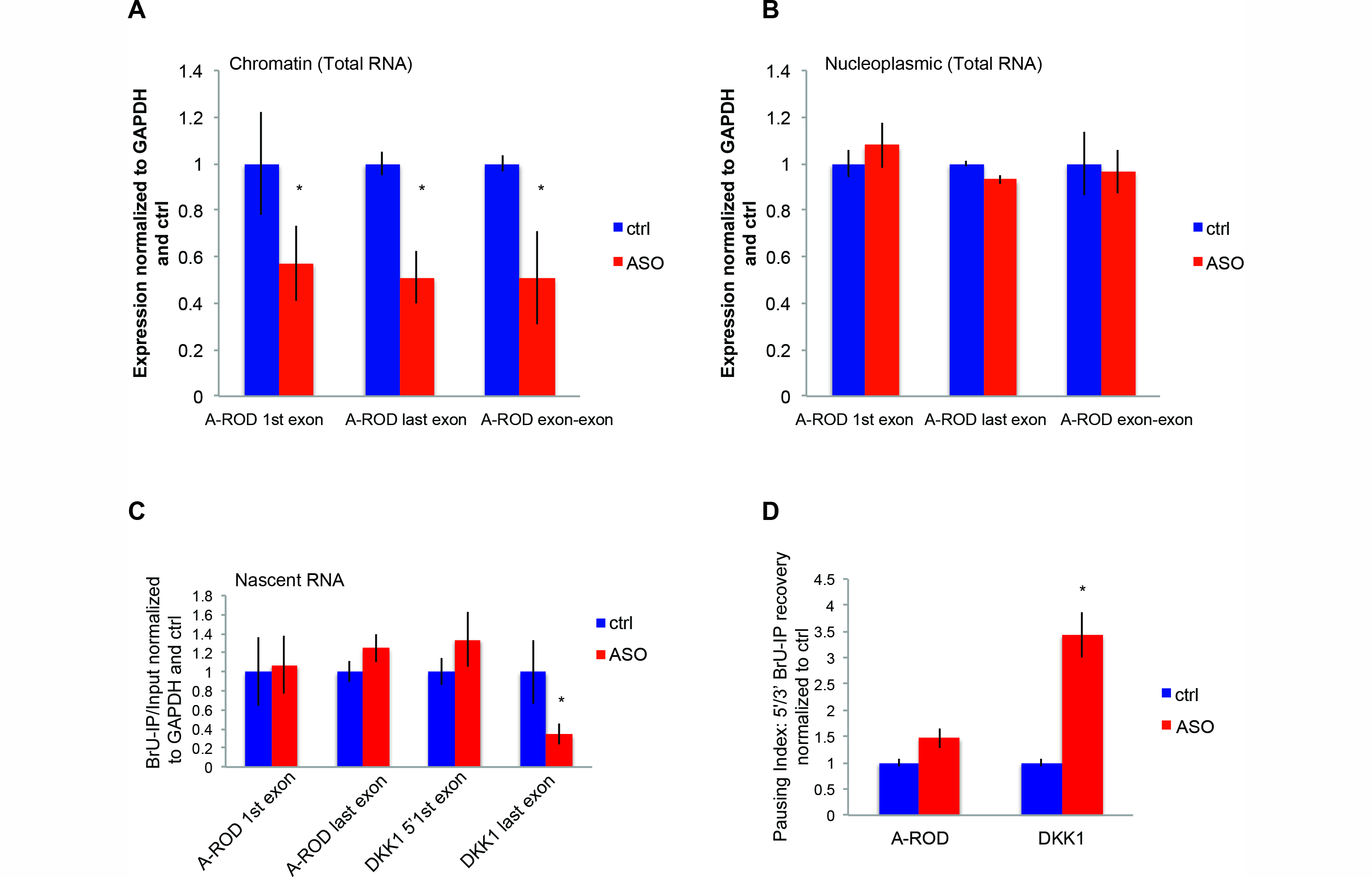
Chromatin-released nascent form of A-ROD enhances DKK1 transcription. (A-B) Expression levels of A-ROD in the steady-state chromatin-associated (A) and nucleoplasmic (B) RNA fraction measured by RT-qPCR in control and A-ROD intronic-ASO treated MCF-7 cells at 16h post transfection. Error bars represent standard deviations from three independent experiments. (C) Normalized RT-qPCR levels in the BrU-labeled nascent RNA (BrU IP done from the chromatin-associated RNA fraction) in control and A-ROD intronic-ASO treated cells. (D) Transcriptional effect assayed by extracting the 5’ to 3’ BrU-IP recovery ratio in control and A-ROD intronic-ASO treated cells.

**Figure 7.**
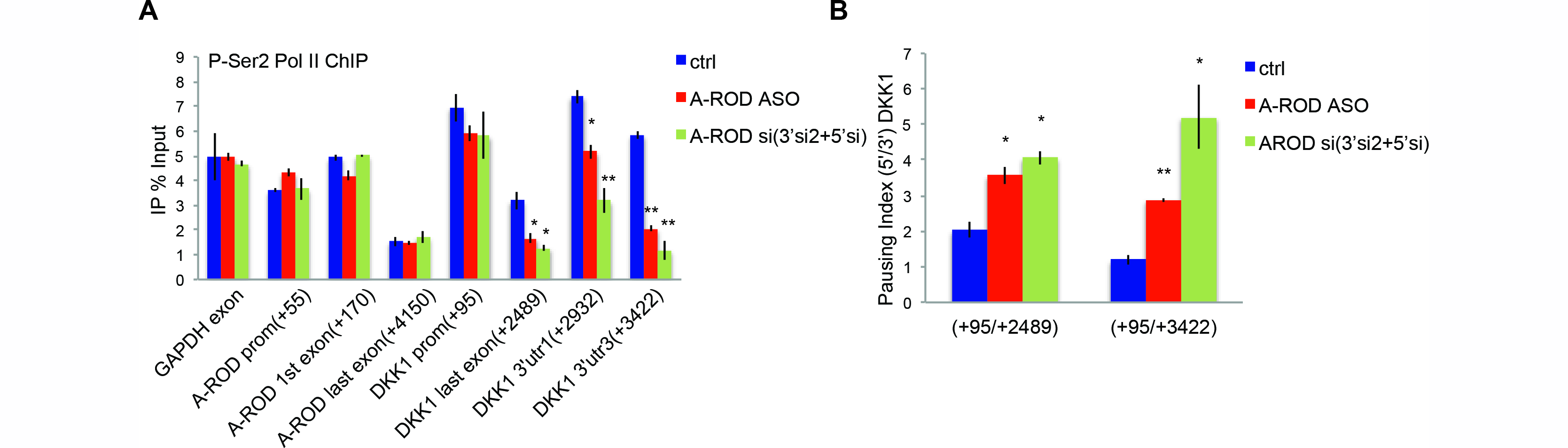
A-ROD enhances DKK1 transcription elongation. (A) ChIP for phospho-Ser2 Pol II in control MCF-7 cells, cells treated with siRNA against A-ROD (here double knock-down transfecting pooled A-ROD.5’si and A-ROD.3’si2) and MCF-7 cells treated with the A-ROD intronic-ASO (here for 24h). Error bars represent standard deviations from three independent experiments. (B) DKK1 transcriptional effect in control, A-ROD siRNA and intronic-ASO treated cells, assayed by extracting the ratio of 5’ to 3’ phospho-Ser2 Pol II ChIP signal depicted in (A).

### siRNA-accessible A-ROD recruits EBP1 to the DKK1 promoter

To determine the molecular mechanism by which A-ROD activates DKK1 transcription we purified endogenous A-ROD RNA using antisense biotinylated oligos tiling the entire A-ROD transcript, modified from previously published methods (Chu et al, 2015; Davis & West, 2015; Vance et al, 2014). Using probes targeting the A-ROD RNA we obtain a specific recovery of A-ROD compared to both control probes and recovery of DKK1 or GAPDH mRNA transcripts (Figure 8A). To assess the binding of A-ROD RNA to the genomic region of DKK1 we analyzed the associated DNA retrieved under the same conditions from the affinity-purified RNA (Figure 8B), showing that A-ROD associates to both the promoter and body of DKK1 but not to the GAPDH promoter as a control locus. In addition, we observe that A-ROD RNA binds to its own genomic locus, which likely reflects its active transcription. To see if A-ROD can recruit specific regulatory proteins to the DKK1 promoter as has been reported for other long ncRNAs and their target genes, we extracted the proteins associated to the affinity-purified A-ROD RNA and identified two candidates by mass spectrometry that show up specifically using the A-ROD antisense probes compared to the LacZ probes; the p48 isoform of EBP1 (ErbB3-binding protein) and HNRNPK (heterogeneous nuclear ribonucleoprotein K) (Figure 8C). Western blot analysis confirms the specific enrichment of EBP1 by A-ROD (Figure 8D) while we could not recapitulate the specificity for HNRNPK although this has been reported to bind specifically to some long ncRNAs (Gumireddy et al, 2013; Huarte et al, 2010) (Figure 8D). The EBP1 isoform p48 has been reported to act as a context-dependent transcriptional activator (Pisapia et al, 2015), and is found to bind RNA in the nucleus (Conrad et al, 2016). To address whether EBP1 can bind to the DKK1 locus and to which extent this is dependent on A-ROD we did ChIP for EBP1 with or without depletion of A-ROD using siRNA (Figure 8E). We show that decreased A-ROD levels leads to decreased EBP1 levels specifically at the DKK1 promoter supporting that the protein is actively recruited to the promoter of DKK1 by A-ROD. Here, GAPDH is used as a control where the promoter is bound by EBP1 but this binding is not affected by A-ROD knock-down. While EBP1 shows a more general expression than A-ROD (Supplementary Figure S4B) the co-expression between DKK1, EBP1 and A-ROD is consistent with a regulatory relationship.

**Figure 8.**
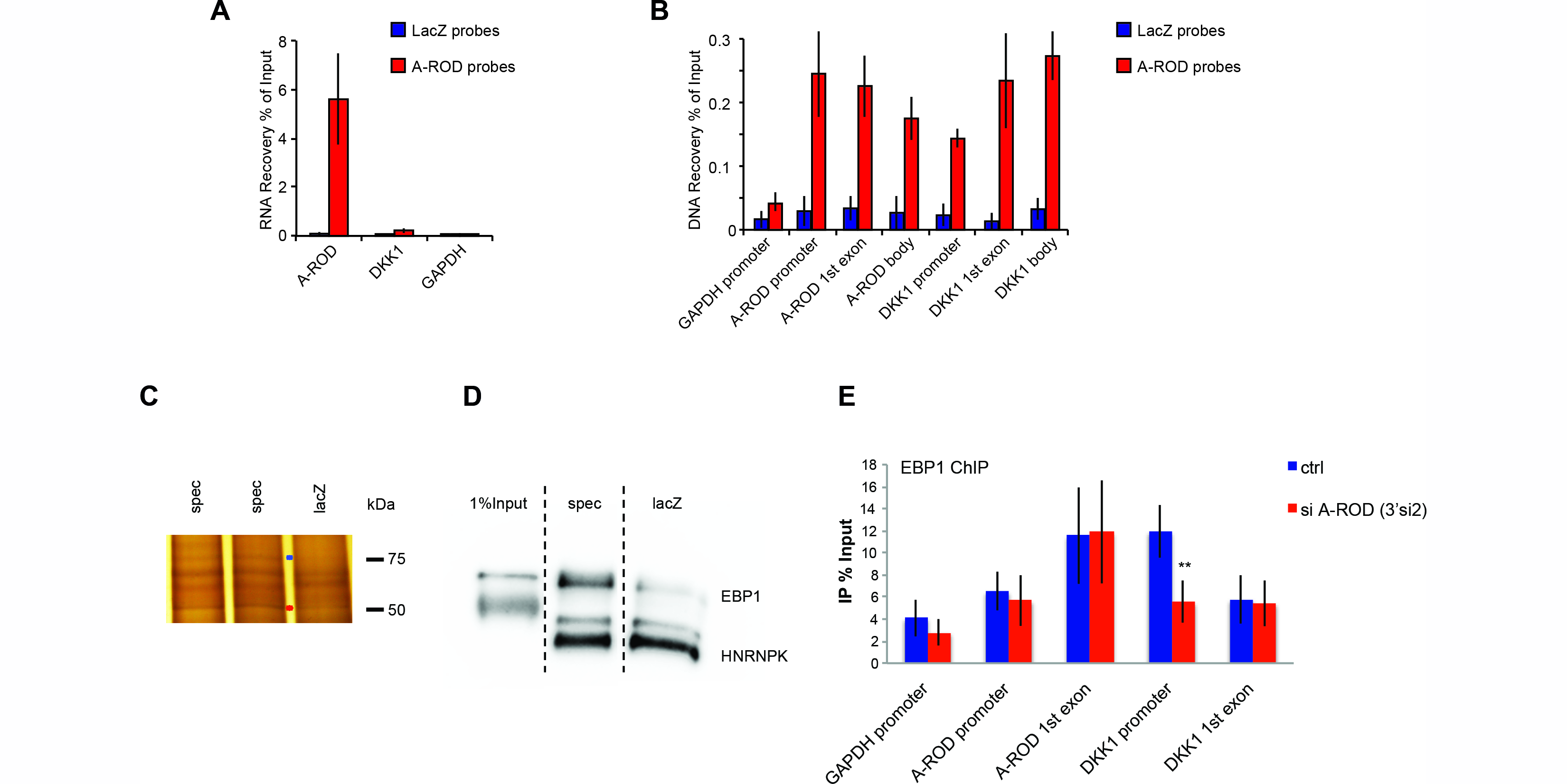
A-ROD recruits EBP1 to DKK1 promoter. (A) RNA recovery of A-ROD, DKK1 or GAPDH transcripts after RNA purification, using biotinylated antisense probes targeting A-ROD (specific) or lacZ (control), measured by RT-qPCR. (B) DNA recovery of the depicted loci measured by qPCR of DNA eluted from RNA purification using biotinylated antisense probes targeting A-ROD or lacZ. Shown are average of at least three independent experiments. (C) Silver staining of SDS-PAGE analyzed proteins, eluted from RNA purification using biotinylated antisense A-ROD specific or lacZ control probes. Marked bands were excised, destained and subjected to mass spectrometry. (D) Western plot for HNRNPK and EBP1 on SDS-PAGE analyzed proteins, eluted from RNA purification using biotinylated antisense A-ROD specific or lacZ control probes. Anti-HNRNPK was first applied on the blot, followed by anti-EBP1. (E) ChIP for EBP1 in control and A-ROD siRNA treated MCF-7 cells. Error bars represent standard deviations from three independent experiments.

Based on the findings reported here, we conclude that the regulatory effects of A-ROD on EBP1 recruitment and DKK1 transcription are dependent on the form of A-ROD RNA that can be targeted by siRNA. We suggest that A-ROD associates with the DKK1 locus and recruits EBP1 to its promoter while loosely associated to or at its release from chromatin, where it gets accessible to siRNA-mediated knock-down, as summarized in the model in Figure 9.

**Figure 9.**
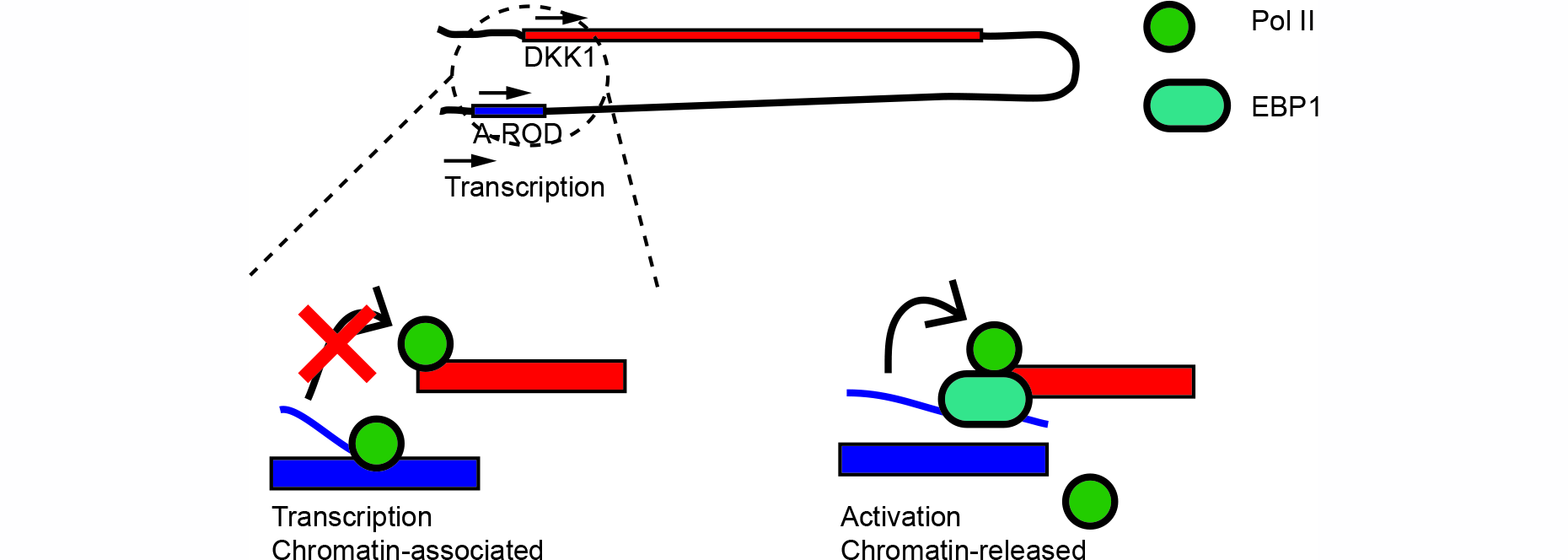
A-ROD ensures transcriptional enhancement of DKK1 in a quasi-cis mode. Model for A-ROD-mediated recruitment of EBP1 and regulation of DKK1 transcription at release from the chromatin-associated template, and within the physical proximity of the pre-established chromosomal interaction.

Our results provide evidence that long ncRNAs interacting with their target genes via pre-established long-range chromosomal interactions are less enriched in chromatin. We propose that this could reflect a general mechanism of long ncRNA function in regulation of target gene expression taking place while loosely associated to or at release from chromatin. We furthermore identify a novel functional interaction between the long ncRNA A-ROD and the EBP1 protein involved in tissue-specific transcription activation of DKK1.

## Discussion

Here, we use a systematic approach to group long ncRNAs by epigenetic marks, showing that around a quarter of long ncRNAs expressed in MCF-7 cells are engaged in strong chromatin interactions as determined by Pol II dependent ChIA-PET. While long ncRNAs are generally chromatin-enriched (Derrien et al, 2012; Werner & Ruthenburg, 2015) those showing strong chromatin interactions are less chromatin-associated. The relatively greater release from chromatin of long ncRNAs in close physical proximity to their target genes suggests that their dissociation from chromatin could play a role in recruiting regulatory protein factors to the promoters of target genes.

We show that A-ROD, a spliced and polyadenylated long ncRNA, can positively regulate the transcription of DKK1 at the level of transcription elongation. It can also recruit the general transcription activator factor EBP1 to the DKK1 promoter via an RNA-dependent mechanism, showing a direct involvement of a long ncRNA in DKK1 transcription in MCF-7 ERa positive breast cancer cells. This could have further implications for gene expression in breast cancer given the correlation in clinical samples of DKK1 and A-ROD (Figure 3D).

How ncRNAs expressed from enhancers can recruit proteins specifically to the promoters of their target genes is a question of high interest to the understanding of regulation of gene expression. While no consensus motifs or sequence complementarity have been found between gene targets and long ncRNAs regulating their expression, the physical proximity is thought to be one of the important factors directing the long ncRNA to its target. How the chromatin interactions are established is not known but based on our and others’ results does not seem to be dependent on the transcription or expression of the long ncRNA. The role of the long ncRNA rather seems to be recruiting transcription factors to chromatin. We propose, based on our results by using siRNA- and ASO-mediated knock-down of nascent A-ROD and analysis in subcellular fractions, that the chromatin-bound form of A-ROD is not the active one required for transcription activation of DKK1. Rather, given that nascent RNA can be targeted by siRNA after a relatively short labeling pulse, the RNA mediates its effect while loosely associated to or at its release from chromatin where the A-ROD locus is already in proximity to the DKK1 locus. The observation that assessment of the siRNA-mediated knock-down of A-ROD when measuring the 3’ end is greater than the 5’ end (Figure 5C) supports that A-ROD has to be fully transcribed in order to be accessible by siRNAs, which implies a functional effect of A-ROD on DKK1 transcription after A-ROD has been transcribed and at its release from chromatin. We therefore suggest a model where transcription of the long ncRNA is not the critical step but rather its release from chromatin where we speculate that it becomes exposed to the nuclear environment and accessible to bind regulatory proteins. While this regulation still occurs on the neighboring gene but requires the release of the long ncRNA from chromatin, it would be in agreement with a quasi-cis mechanism of action where the targeting specificity is mediated by the chromatin interaction established prior to and independent of the expression of the long ncRNA, and the regulatory effect is mediated by the long ncRNA at release from chromatin.

Long ncRNAs expressed from enhancers are often polyadenylated and more stable than eRNAs that are mostly non-polyadenylated and short-lived and expected not to dissociate from chromatin (Andersson et al, 2014; Kim et al, 2010; Wang et al, 2011a). While eRNAs have been shown to increase the affinity for DNA-binding proteins to their site of transcription (Sigova et al, 2015) the active recruitment of proteins by spliced long ncRNA at release from chromatin could be one of the factors distinguishing these two groups of non-coding transcripts expressed from enhancers. The involvement of chromatin-release of long ncRNAs, as an exerted aspect of their functionality, is a novel finding that should be explored further to establish its general importance and impact in targeted gene regulation and long ncRNA and enhancer function.

## Materials and Methods

Detailed Methods can be found in the Supplementary Materials and Methods.

## ChIA-PET

ChIA-PET reads in MCF-7 from 4 replicates (Li et al., Cell 2012; ENCODE 2012) were pooled and filtered for a cutoff score of 200. Pairs of interacting long ncRNA with annotated ENCODE V22 genes were identified by intersecting intervals of □2 kb around the TSS with the ChIA-PET nodes, using bedtools (Quinlann and Hall, 2010).

## RNA-seq and expression analysis

Chromatin associated RNA-seq libraries in MCF-7 cells were prepared as in (Conrad et al, 2014) and mapping of strand-specific reads was done with the STAR algorithm (Dobin et al, 2013). Nucleoplasmic-enriched RNA-seq libraries were prepared from the nucleoplasmic fraction upon cell fractionation as in (Conrad et al, 2014) and mapping was done using STAR. Differential expression analysis of chromatin-associated versus nucleoplasmic RNA as well of available GRO-seq data (Hah et al, 2013) (GEO accession number GSE43835) was performed using DESeq2 (Love et al, 2014). Expression specificity was extracted using the same script for extracting CpG hypo-methylation specificity (see Supplementary Methods under ‘Methylation status analysis’), loading with strand-specific read counts from RNA-seq data (ENCODE, 2012).

## Set of long ncRNAs

Long ncRNAs used in this study are a combination of ENCODE V22 annotated long ncRNAs, previously annotated long ncRNAs (Cabili et al, 2011) and *de novo* transcript assembly using MCF-7 chromatin-associated RNA-seq data (GEO accession number GSE69507) (see also *‘De novo* transcript assembly’ in Supplementary Methods).

## K-means clustering

Peaks from available MCF-7 histone mark ChIP-seq data (ENCODE H3K4me3 and H3K27ac and H3K4me1 from (Joseph et al, 2010) (GEO GSM588569) were called at p < 0.01 using MACS2 (version 2.0.10.20120913) (Zhang et al, 2008). Signal values from long ncRNA overlapping peaks were summed across transcript unit length. All data values, including *log* transformed GRO-seq and CHR-RNA-seq RPKM, were rescaled in the range between 0 and 1 using min-max normalization. K-means clustering was performed with R function *kmeans*, with k = 15 (at this number the total within-cluster sum of squares reached a minimum plateau) and nstart = 100.

## Linear Regression and PCA

Linear regression was performed with R function lm on Z-score standardized values of the included parameters for long ncRNAs with more than 10 GRO-seq reads per kb. PCA was performed with R function prcomp and visualized with ggbiplot (https://github.com/vqv/ggbiplot).

## Breast cancer RNA-seq data

RNA-seq data from ER positive and ER negative breast cancer samples were downloaded from the Cancer Genome Atlas (TCGA; TCGA_BRCA_exp_HiSeqV2_exon_2014-08-28). Total exon RPKM were summed per transcript.

## GTEx Project data

Expression data from the Genotype-Tissue Expression (GTEx) Project were obtained from the GTEx portal, latest release version V6p.

## Purification of nascent RNA

MCF-7 cells in culture, seeded one day before, were incubated with 2.5 mM final concentration of 5-bromouridine for 15 min, then harvested on ice and subjected to cell fractionation according to (Conrad and Ørom, Methods Mol Biol 2017). 30 μl of Protein G Dynabeads were incubated with 5 μg of purified mouse anti-BrdU antibody (BD Biosciences) for 1 h at room temperature. ~4 μg of BrU-labeled chromatin-associated or nucleoplasmic RNA were incubated with precoupled antibody (1 h at room temperature in the presence of RNase inhibitors) followed by five washes in ice cold PBS. Nascent RNA was eluted twice from the beads with competitive elution (25 mM 5-bromouridine in PBS) for 1 h at 4 □C. Eluates were pooled and precipitated with EtOH or using concentrator columns (Zymo Research). All of the eluted nascent RNA (and ~400 ng of chromatin-associated or nucleoplasmic input RNA) was subjected to reverse transcription with random hexamer primers.

## Purification of target RNA and associated DNA and proteins

Purification of target RNA and associated DNA and proteins was done based on the previously published methods ChIRP (Chu et al, 2015) and CHART (Davis & West, 2015) with modifications (see Supplementary Methods).

## Cell Culture and knock-down

MCF-7 cells were grown in DMEM (10% FCS) or in hormone-free medium (high glucose DMEM without phenol red) supplemented with 5 % charcoal stripped FCS for 72 h before E2 induction. For siRNA mediated knockdown cells were transfected twice (48 h interval) with a 35 nM final concentration of siRNA. Cells were harvested 84-96 h after seeding. Biological replicates with at least two different siRNA sequences were performed. As a negative control, cells were transfected with 35 nM of the NC1 duplex (Integrated DNA Technologies, Inc). For ASO knock-down of A-ROD, cells were transfected with a 50 nM final concentration of the intronic ASO and harvested after 16 h. As a negative control cells were transfected with 50 nM of pooled lacZ oligos. SiRNA and ASO sequences are listed in the Supplementary Materials and Methods.

## Chromosome conformation capture (3C)

3C was performed as previously described (Lai et al., Nature 2013) with minor modifications (see Supplementary Methods).

## Statistical analysis

P-values were extracted using Student’s two-tailed t-test for all features with normal distribution, otherwise Wilcoxon-Mann-Whitney test was applied.

## Data access

Data from chromatin-associated and nucleoplasmic RNA sequencing and DNA methylation bisulfite sequencing have been deposited to GEO with accession number GSE69507.

## Acknowledgments

We thank Melissa J. Fullwood for critical reading of the manuscript. E.N. has been funded by a postdoctoral fellowship from the Alexander von Humboldt Foundation. Work in the authors’ laboratory is funded by the Alexander von Humboldt Foundation and the German ministry for research and education through the Sofja Kovalevskaja Award to U.A.Ø.

## Author Contributions

E.N. performed experiments and bioinformatic analysis, apart from the DNA hypomethylation specificity code written by J.M. J.L. performed experiments. A.M. and J.M. supervised bioinformatic analysis. E.N. and U.A.Ø. conceived experiments, interpreted data and wrote the manuscript. U.A.Ø. supervised research. All authors read and approved the manuscript.

## Supplementary Figure Legends

**Supplementary Figure S1.**
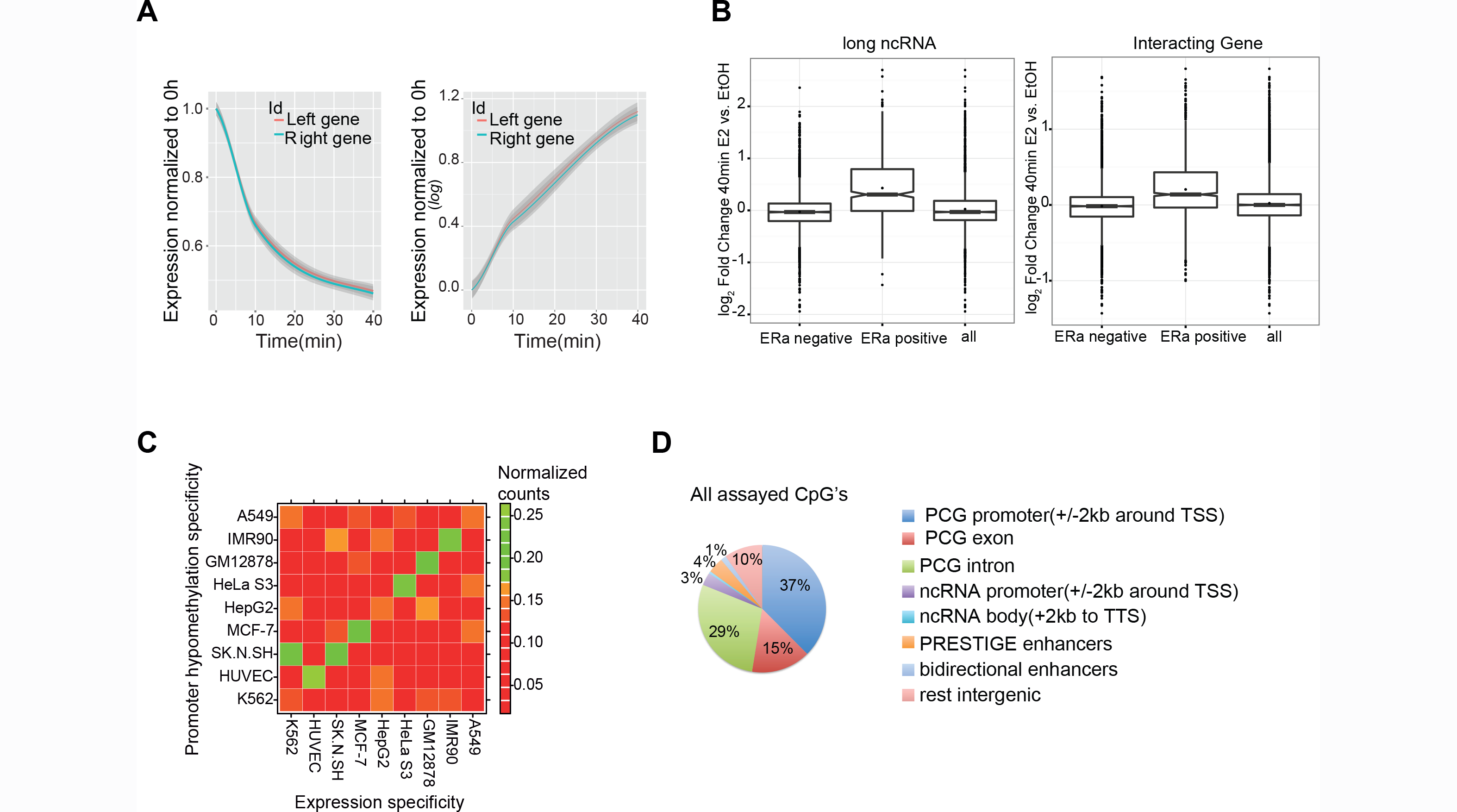
Long ncRNA expression and epigenetic marks characteristics. (A) E2-mediated time response of downregulated (left panel) and upregulated (right panel) randomly selected ChIA-PET interacting gene-gene pairs (with a ChIA-PET score > 200). GRO-seq RPKM from 10 and 40 min of E2 treatment were normalized to 0 h (EtOH control). In the right panel y-axis is in the *log* scale. Fitted regression model lines (loess curves) are shown and the 95% confidence interval is grey shaded. The analysis was ran several times (> 10) with different pools of randomly sampled ChIA-PET interconnected gene-gene pairs, producing similarly overlapping time responses. (B) Boxplot showing *DESeq2* generated regularized log2 fold changes in expression of 40 min E2 treated MCF-7 cells versus control, using GRO-seq, for long ncRNAs (left) and their ChIA-PET identified interacting genes (right), sub-categorized as either ‘ERa positive’ or ‘ERa negative’, based on the presence or not of ERa binding sites (ChIP-seq peaks) +/-10 kb around the transcription start site (TSS). For the 4,467 long ncRNAs (left): 513 have ERa +/-10 kb around the TSS (‘ERa positive’), and of those, 382 are E2-upregulated showing significantly higher log2 fold changes in expression than the rest (‘ERa negative’ or compared to all; Wilcoxon-Mann-Whitney test p-value < 2.2e-16). In addition, ERa-positive long ncRNAs are significantly enriched amongst E2-upregulated cases (Fisher’s exact test p-value = 3.237e-11, odds ratio = 1.63). For the 7,272 interacting genes (right): 1238 are ERa-positive and of those, 860 are E2-upregulated showing significantly higher log2 fold changes in expression (Wilcoxon-Mann-Whitney test p-value < 2.2e-16). In addition, ERa-positive genes are significantly enriched amongst E2-upregulated cases (Fisher’s exact test p-value = 2.457e-11, odds ratio 1.39). (C) Heatmap showing correspondence (normalized counts; observed/expected counts for each cell-type combination) between long ncRNA expression specificity (extracted from ENCODE available nuclear polyA+ RNA-seq data) and associated promoter hypomethylation cell-type specificity (extracted from Methyl 450K Array b-values). (D) Pie chart of the genomic distribution of SureSelect assayed CpG’s common for 0 h (EtOH control) and 40 min estradiol (E2) treatment; in total 3,525,513 CpG’s with an applied read coverage cutoff of 5. Protein coding gene (PCG) and long ncRNA promoters are here defined as ±2kb intervals around the transcription start site (TSS). PRESTIGE enhancers are from (Corradin et al., 2014). Putative bidirectional enhancers (±1kb extended) are from (Andersson et al., 2014).

**Supplementary Figure S2.**
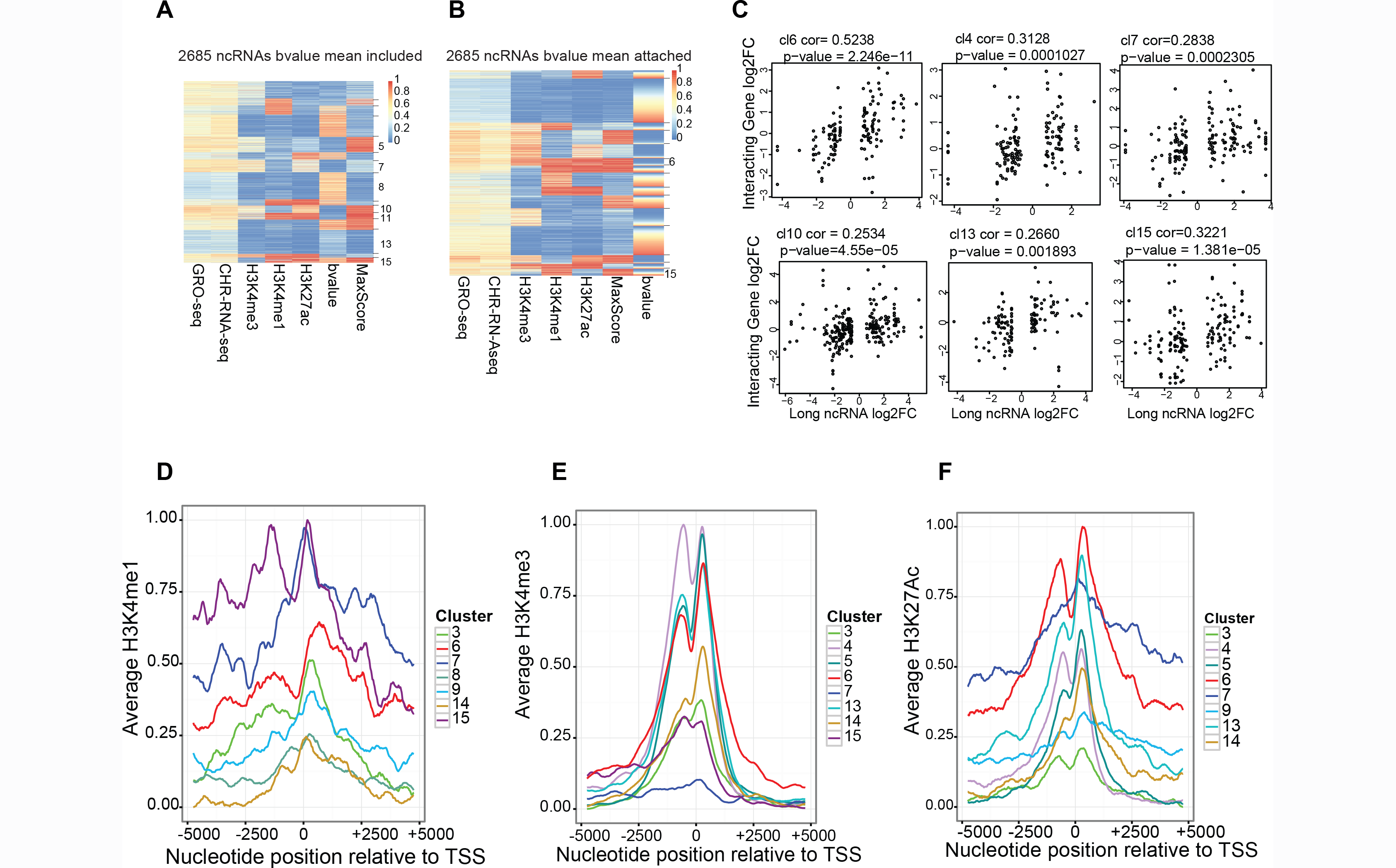
Grouping of long ncRNAs based on expression and epigenetic marks. (A) K-means clustering of 2,685 long ncRNAs including expression (GRO-seq and chromatin-associated RNA-seq), histone mark signal values, ChIA-PET maximum interaction scores and SureSelect assayed DNA methylation (promoter b-value mean). (B) K-means clustering was initially done for 4,467 long ncRNAs without DNA methylation as a clustering parameter (Figure 1D) and then the promoter b-value mean was attached for the entries with assayed DNA methylation (2,685) producing the heatmap shown here (entries have been ordered within each cluster according to increasing DNA methylation). (C) Scatterplots showing correlation of E2-mediated log2 fold changes in expression (40 min E2 treatment vs. EtOH control; GRO-seq) for long ncRNAs of clusters 6, 4, 7, 10, 13, 15 (Figure 1D) and ChIA-PET identified interacting genes. (D-F) Positional distribution of histone marks around the transcription start site (TSS) of the clustered long ncRNAs. Average histone mark signal (IP/Input) for (D) H3K4me1, (E) H3K4me3 and (F) H3K27Ac was smoothened over a 500bp sliding window average (sliding step 1) ±5kb around the TSS of the clustered long ncRNAs followed by min-max rescaling for all.

**Supplementary Figure S3.**
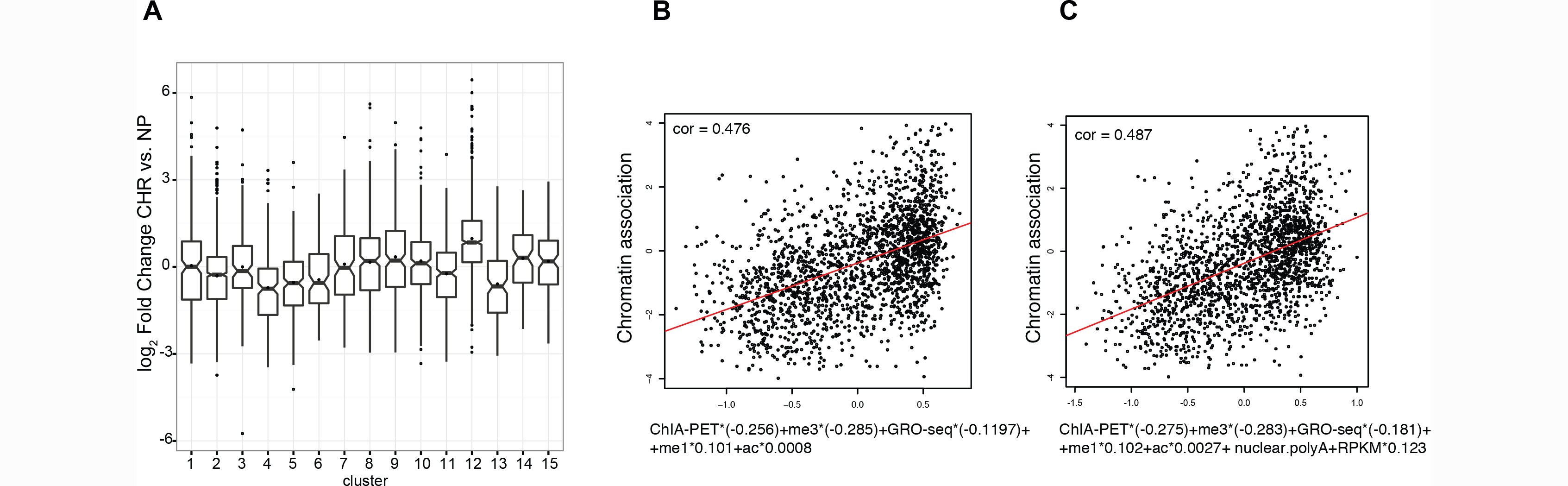
Chromatin-association of long ncRNAs. (A) DESeq2 generated log2 fold changes in expression using total chromatin-associated versus nucleoplasmic RNA-seq, for the clustered long ncRNAs of Figure 1D. (See also Figure 2A–B). Significantly nucleoplasmic-enriched long ncRNAs (at *DESeq2* p-adjusted < 0.1) are significantly enriched in clusters 4, 5, 6 and 13 (Fisher’s exact test p-value = [5.676e-12, 1.026e-12, 2.653e-06, 1.089e-07] and odds ratio = [2.53, 2.45, 2.35 and 2.36], respectively). (B) Linear regression analysis by incorporating the parameters ChIA-PET interaction score, GRO-seq RPKM, H3K4me3, H3K4me1 and H3K27ac, in the prediction of chromatin-association of long ncRNAs. The generated numeric coefficients for each variable are included in the type below the scatterplot. The red line represents the linear regression model fit (correlation = 0.476, p-value < 2.2e-16). (C) Same as in B but with one additional variable included, that is nuclear polyA+ RPKM.

**Supplementary Figure S4.**
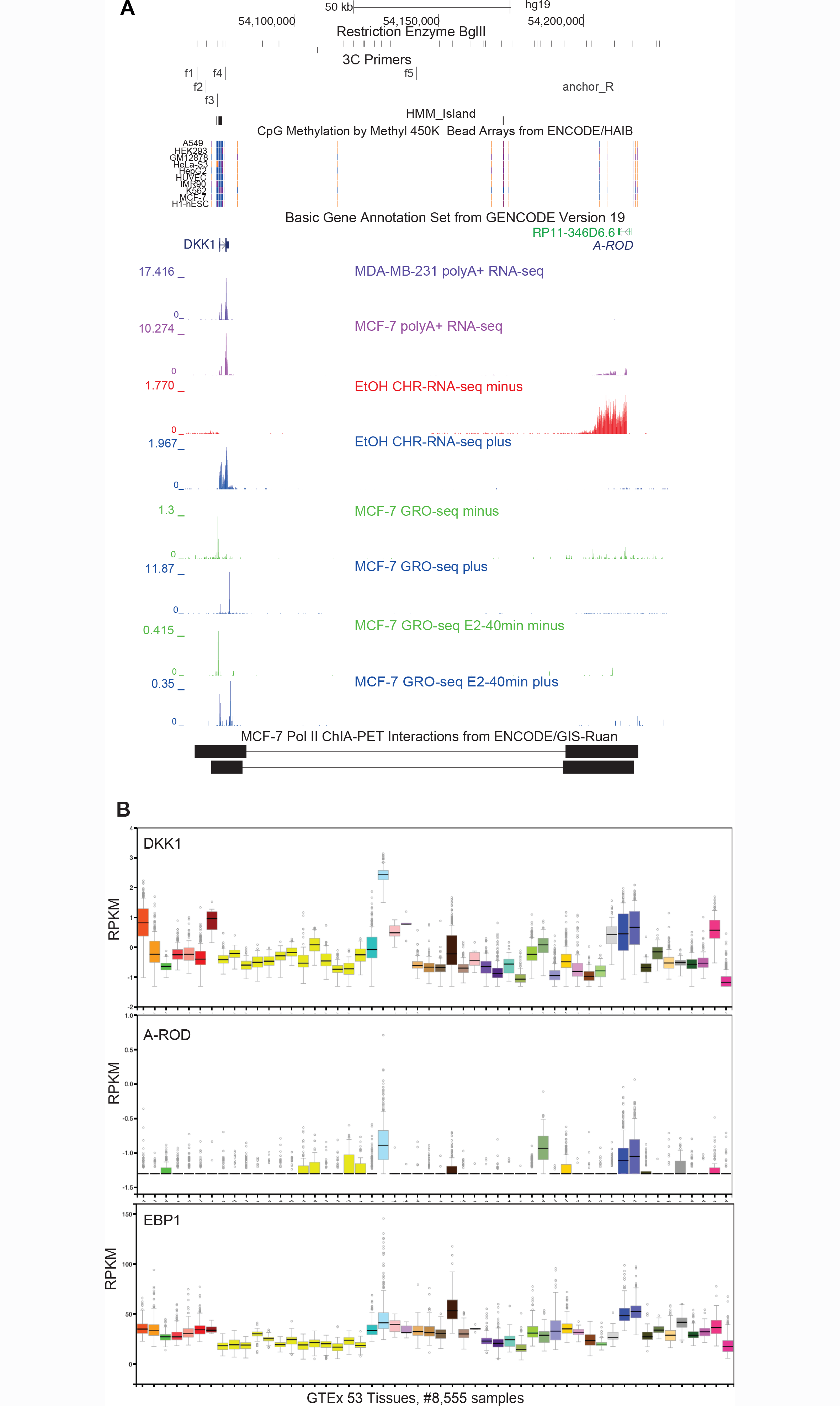
A-ROD as a tissue-specific activator of DKK1 expression. (A) UCSC genome browser overview of the DKK1 and A-ROD (RP11-346D6.6) genomic loci. Location of the 3C primers, polyA+ RNA-seq data from MDA-MB-231 and MCF-7 (GEO GSE37918), chromatin-associated RNA-seq in MCF-7 and GRO-seq (GEO GSE43835) are shown (*y* axis reads per million). (B) Expression of DKK1, A-ROD and EBP1 (RPKM; plotted in the *log* scale for DKK1 and A-ROD) taken from the GTEx Project (version V6p).

**Supplementary Figure S5.**
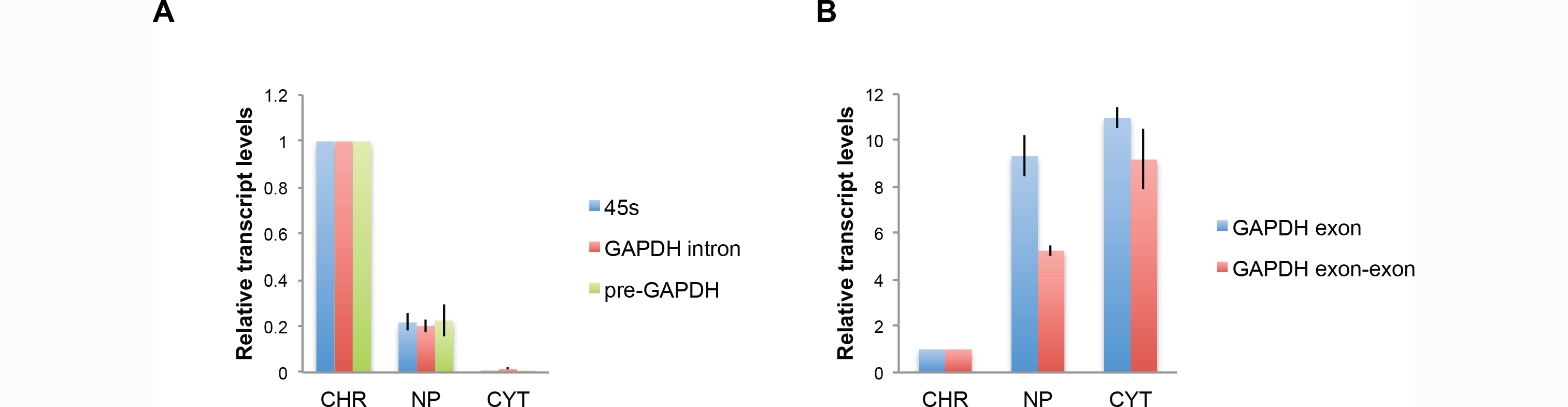
Cell fractionation control. (A-B) RT-qPCR assaying the relative transcript levels of relevant markers in the three cellular fractions, normalized to the respective values in the chromatin-associated fraction. Error bars represent standard deviations from three independent fractionation experiments.

**Supplementary Figure S6.**
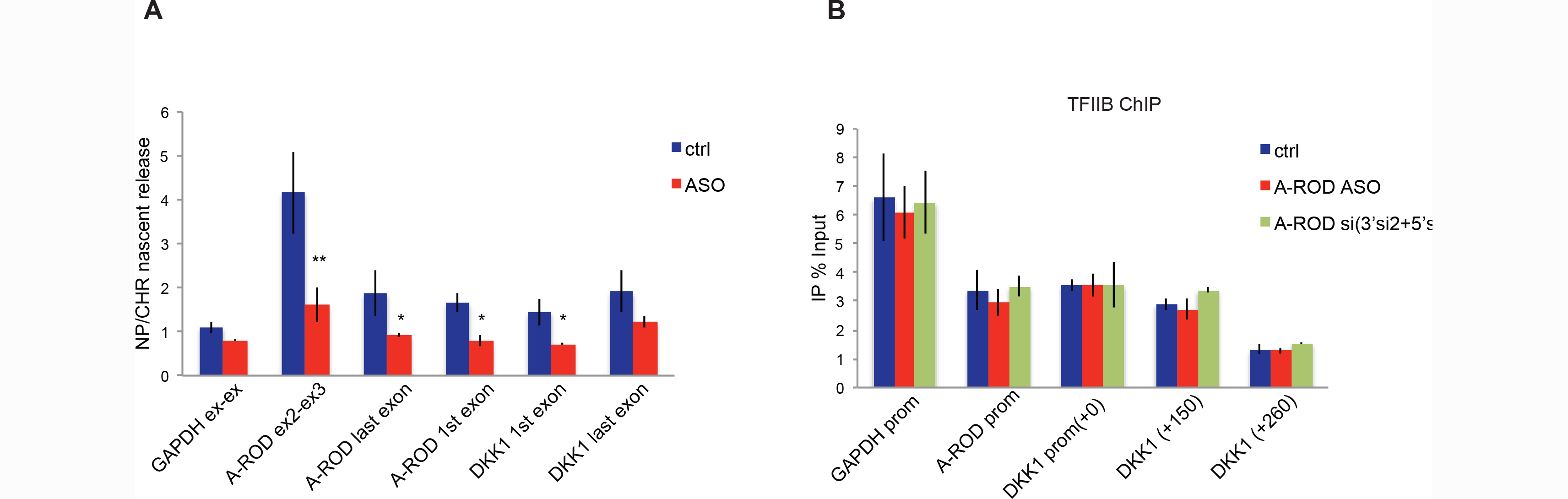
A-ROD regulates DKK1 transcription at its release from chromatin. (A) Nascent transcript release assayed by extracting the ratio of BrU-labeled nucleoplasmic to BrU-labeled chromatin-associated RNA. Error bars represent standard deviations from three independent experiments. (B) TFIIB ChIP in control and A-ROD siRNA (here pooled A-ROD.5’si and A-ROD.3’si2) or A-ROD intronic-ASO treated cells. Error bars represent standard deviations from three independent experiments.

## Supplementary Tables

**Supplementary Table S1. The 4,467 long ncRNAs expressed in MCF-7 cells.**

**Supplementary Table S2. Long ncRNAs enriched at chromatin.**

**Supplementary Table S3. Long ncRNAs enriched in the nucleoplasm.**

## References

Adelman K, Lis JT (2012) Promoter-proximal pausing of RNA polymerase II: emerging roles in metazoans. Nat Rev Genet 13: 720–731

Andersson R, Gebhard C, Miguel-Escalada I, Hoof I, Bornholdt J, Boyd M, Chen Y, Zhao X, Schmidl C, Suzuki T, Ntini E, Arner E, Valen E, Li K, Schwarzfischer L, Glatz D, Raithel J, Lilje B, Rapin N, Bagger FO, Jorgensen M, Andersen PR, Bertin N, Rackham O, Burroughs AM, Baillie JK, Ishizu Y, Shimizu Y, Furuhata E, Maeda S, Negishi Y, Mungall CJ, Meehan TF, Lassmann T, Itoh M, Kawaji H, Kondo N, Kawai J, Lennartsson A, Daub CO, Heutink P, Hume DA, Jensen TH, Suzuki H, Hayashizaki Y, Muller F, Forrest AR, Carninci P, Rehli M, Sandelin A (2014) An atlas of active enhancers across human cell types and tissues. Nature 507: 455–461

Arner E, Daub CO, Vitting-Seerup K, Andersson R, Lilje B, Drablos F, Lennartsson A, Ronnerblad M, Hrydziuszko O, Vitezic M, Freeman TC, Alhendi AM, Arner P, Axton R, Baillie JK, Beckhouse A, Bodega B, Briggs J, Brombacher F, Davis M, Detmar M, Ehrlund A, Endoh M, Eslami A, Fagiolini M, Fairbairn L, Faulkner GJ, Ferrai C, Fisher ME, Forrester L, Goldowitz D, Guler R, Ha T, Hara M, Herlyn M, Ikawa T, Kai C, Kawamoto H, Khachigian LM, Klinken SP, Kojima S, Koseki H, Klein S, Mejhert N, Miyaguchi K, Mizuno Y, Morimoto M, Morris KJ, Mummery C, Nakachi Y, Ogishima S, Okada-Hatakeyama M, Okazaki Y, Orlando V, Ovchinnikov D, Passier R, Patrikakis M, Pombo A, Qin XY, Roy S, Sato H, Savvi S, Saxena A, Schwegmann A, Sugiyama D, Swoboda R, Tanaka H, Tomoiu A, Winteringham LN, Wolvetang E, Yanagi-Mizuochi C, Yoneda M, Zabierowski S, Zhang P, Abugessaisa I, Bertin N, Diehl AD, Fukuda S, Furuno M, Harshbarger J, Hasegawa A, Hori F, Ishikawa-Kato S, Ishizu Y, Itoh M, Kawashima T, Kojima M, Kondo N, Lizio M, Meehan TF, Mungall CJ, Murata M, Nishiyori-Sueki H, Sahin S, Nagao-Sato S, Severin J, de Hoon MJ, Kawai J, Kasukawa T, Lassmann T, Suzuki H, Kawaji H, Summers KM, Wells C, Hume DA, Forrest AR, Sandelin A, Carninci P, Hayashizaki Y (2015) Transcribed enhancers lead waves of coordinated transcription in transitioning mammalian cells. Science 347: 1010–1014

Bulger M, Groudine M (2011) Functional and mechanistic diversity of distal transcription enhancers. Cell 144: 327–339

Cabili MN, Trapnell C, Goff L, Koziol M, Tazon-Vega B, Regev A, Rinn JL (2011) Integrative annotation of human large intergenic noncoding RNAs reveals global properties and specific subclasses. Genes Dev 25: 1915–1927

Chu C, Zhang QC, da Rocha ST, Flynn RA, Bharadwaj M, Calabrese JM, Magnuson T, Heard E, Chang HY (2015) Systematic discovery of Xist RNA binding proteins. Cell 161: 404–416

Conrad T, Albrecht AS, de Melo Costa VR, Sauer S, Meierhofer D, Orom UA (2016) Serial interactome capture of the human cell nucleus. Nat Commun 7: 11212

Conrad T, Marsico A, Gehre M, Orom UA (2014) Microprocessor activity controls differential miRNA biogenesis In Vivo. Cell Rep 9: 542–554

Core LJ, Martins AL, Danko CG, Waters CT, Siepel A, Lis JT (2014) Analysis of nascent RNA identifies a unified architecture of initiation regions at mammalian promoters and enhancers. Nat Genet 46: 1311–1320

Core LJ, Waterfall JJ, Lis JT (2008) Nascent RNA sequencing reveals widespread pausing and divergent initiation at human promoters. Science 322: 1845–1848

Davis CP, West JA (2015) Purification of specific chromatin regions using oligonucleotides: capture hybridization analysis of RNA targets (CHART). Methods Mol Biol 1262: 167–182

De Santa F, Barozzi I, Mietton F, Ghisletti S, Polletti S, Tusi BK, Muller H, Ragoussis J, Wei CL, Natoli G (2010) A large fraction of extragenic RNA pol II transcription sites overlap enhancers. PLoS Biol 8: e1000384

Derrien T, Johnson R, Bussotti G, Tanzer A, Djebali S, Tilgner H, Guernec G, Martin D, Merkel A, Knowles DG, Lagarde J, Veeravalli L, Ruan X, Ruan Y, Lassmann T, Carninci P, Brown JB, Lipovich L, Gonzalez JM, Thomas M, Davis CA, Shiekhattar R, Gingeras TR, Hubbard TJ, Notredame C, Harrow J, Guigo R (2012) The GENCODE v7 catalog of human long noncoding RNAs: Analysis of their gene structure, evolution, and expression. Genome Res 22: 1775–1789

Dobin A, Davis CA, Schlesinger F, Drenkow J, Zaleski C, Jha S, Batut P, Chaisson M, Gingeras TR (2013) STAR: ultrafast universal RNA-seq aligner. Bioinformatics 29: 15–21

ENCODE (2012) An integrated encyclopedia of DNA elements in the human genome. Nature 489: 57–74

Gagnon KT, Li L, Chu Y, Janowski BA, Corey DR (2014) RNAi factors are present and active in human cell nuclei. Cell Rep 6: 211–221

Gomez JA, Wapinski OL, Yang YW, Bureau JF, Gopinath S, Monack DM, Chang HY, Brahic M, Kirkegaard K (2013) The NeST long ncRNA controls microbial susceptibility and epigenetic activation of the interferon-gamma locus. Cell 152: 743–754

Gumireddy K, Li A, Yan J, Setoyama T, Johannes GJ, Orom UA, Tchou J, Liu Q, Zhang L, Speicher DW, Calin GA, Huang Q (2013) Identification of a long non-coding RNA-associated RNP complex regulating metastasis at the translational step. Embo J 32: 2672–2684

Hah N, Danko CG, Core L, Waterfall JJ, Siepel A, Lis JT, Kraus WL (2011) A rapid, extensive, and transient transcriptional response to estrogen signaling in breast cancer cells. Cell 145: 622–634

Hah N, Murakami S, Nagari A, Danko CG, Kraus WL (2013) Enhancer transcripts mark active estrogen receptor binding sites. Genome Res

Heidari N, Phanstiel DH, He C, Grubert F, Jahanbani F, Kasowski M, Zhang MQ, Snyder MP (2014) Genome-wide map of regulatory interactions in the human genome. Genome Res 24: 19051917

Hsieh CL, Fei T, Chen Y, Li T, Gao Y, Wang X, Sun T, Sweeney CJ, Lee GS, Chen S, Balk SP, Liu XS, Brown M, Kantoff PW (2014) Enhancer RNAs participate in androgen receptor-driven looping that selectively enhances gene activation. Proc Natl Acad Sci U S A 111: 7319–7324

Huarte M, Guttman M, Feldser D, Garber M, Koziol MJ, Kenzelmann-Broz D, Khalil AM, Zuk O, Amit I, Rabani M, Attardi LD, Regev A, Lander ES, Jacks T, Rinn JL (2010) A large intergenic noncoding RNA induced by p53 mediates global gene repression in the p53 response. Cell 142: 409–419

Joseph R, Orlov YL, Huss M, Sun W, Kong SL, Ukil L, Pan YF, Li G, Lim M, Thomsen JS, Ruan Y, Clarke ND, Prabhakar S, Cheung E, Liu ET (2010) Integrative model of genomic factors for determining binding site selection by estrogen receptor-alpha. Mol Syst Biol 6: 456

Khalil AM, Guttman M, Huarte M, Garber M, Raj A, Rivea Morales D, Thomas K, Presser A, Bernstein BE, van Oudenaarden A, Regev A, Lander ES, Rinn JL (2009) Many human large intergenic noncoding RNAs associate with chromatin-modifying complexes and affect gene expression. Proc Natl Acad Sci U S A 106: 11667–11672

Kim TK, Hemberg M, Gray JM, Costa AM, Bear DM, Wu J, Harmin DA, Laptewicz M, Barbara-Haley K, Kuersten S, Markenscoff-Papadimitriou E, Kuhl D, Bito H, Worley PF, Kreiman G, Greenberg ME (2010) Widespread transcription at neuronal activity-regulated enhancers. Nature 465: 182–187

Kim YW, Lee S, Yun J, Kim A (2015) Chromatin looping and eRNA transcription precede the transcriptional activation of gene in the beta-globin locus. Biosci Rep 35

Lai F, Orom UA, Cesaroni M, Beringer M, Taatjes DJ, Blobel GA, Shiekhattar R (2013) Activating RNAs associate with Mediator to enhance chromatin architecture and transcription. Nature 494: 497–501

Lam MT, Li W, Rosenfeld MG, Glass CK (2014) Enhancer RNAs and regulated transcriptional programs. Trends Biochem Sci 39: 170–182

Li G, Ruan X, Auerbach RK, Sandhu KS, Zheng M, Wang P, Poh HM, Goh Y, Lim J, Zhang J, Sim HS, Peh SQ, Mulawadi FH, Ong CT, Orlov YL, Hong S, Zhang Z, Landt S, Raha D, Euskirchen G, Wei CL, Ge W, Wang H, Davis C, Fisher-Aylor KI, Mortazavi A, Gerstein M, Gingeras T, Wold B, Sun Y, Fullwood MJ, Cheung E, Liu E, Sung WK, Snyder M, Ruan Y (2012) Extensive promoter-centered chromatin interactions provide a topological basis for transcription regulation. Cell 148: 84–98

Li W, Notani D, Rosenfeld MG (2016) Enhancers as non-coding RNA transcription units: recent insights and future perspectives. Nature Reviews Genetics 17: 207–223

Love MI, Huber W, Anders S (2014) Moderated estimation of fold change and dispersion for RNA-seq data with DESeq2. Genome Biol 15: 550

Marchese FP, Grossi E, Marin-Bejar O, Bharti SK, Raimondi I, Gonzalez J, Martinez-Herrera DJ, Athie A, Amadoz A, Brosh RM, Jr., Huarte M (2016) A Long Noncoding RNA Regulates Sister Chromatid Cohesion. Mol Cell

Mikheev AM, Mikheeva SA, Maxwell JP, Rivo JV, Rostomily R, Swisshelm K, Zarbl H (2008) Dickkopf-1 mediated tumor suppression in human breast carcinoma cells. Breast Cancer Res Treat 112: 263–273

Niida A, Hiroko T, Kasai M, Furukawa Y, Nakamura Y, Suzuki Y, Sugano S, Akiyama T (2004) DKK1, a negative regulator of Wnt signaling, is a target of the beta-catenin/TCF pathway. Oncogene 23: 8520–8526

Orom UA, Derrien T, Beringer M, Gumireddy K, Gardini A, Bussotti G, Lai F, Zytnicki M, Notredame C, Huang Q, Guigo R, Shiekhattar R (2010) Long noncoding RNAs with enhancer-like function in human cells. Cell 143: 46–58

Orom UA, Shiekhattar R (2013) Long noncoding RNAs usher in a new era in the biology of enhancers. Cell 154: 1190–1193

Pekowska A, Benoukraf T, Zacarias-Cabeza J, Belhocine M, Koch F, Holota H, Imbert J, Andrau JC, Ferrier P, Spicuglia S (2011) H3K4 tri-methylation provides an epigenetic signature of active enhancers. Embo J 30: 4198–4210

Pisapia L, Barba P, Cortese A, Cicatiello V, Morelli F, Del Pozzo G (2015) EBP1 protein modulates the expression of human MHC class II molecules in non-hematopoietic cancer cells. Int J Oncol 47: 481–489

Qiao L, Xu ZL, Zhao TJ, Ye LH, Zhang XD (2008) Dkk-1 secreted by mesenchymal stem cells inhibits growth of breast cancer cells via depression of Wnt signalling. Cancer Lett 269: 67–77

Quinn JJ, Chang HY (2016) Unique features of long non-coding RNA biogenesis and function. Nat Rev Genet 17: 47–62

Schaukowitch K, Joo JY, Liu X, Watts JK, Martinez C, Kim TK (2014) Enhancer RNA Facilitates NELF Release from Immediate Early Genes. Mol Cell 56: 29–42

Sigova AA, Abraham BJ, Ji X, Molinie B, Hannett NM, Guo YE, Jangi M, Giallourakis CC, Sharp PA, Young RA (2015) Transcription factor trapping by RNA in gene regulatory elements. Science 350: 978–981

Trimarchi T, Bilal E, Ntziachristos P, Fabbri G, Dalla-Favera R, Tsirigos A, Aifantis I (2014) Genome-wide Mapping and Characterization of Notch-Regulated Long Noncoding RNAs in Acute Leukemia. Cell 158: 593–606

Vance KW, Sansom SN, Lee S, Chalei V, Kong L, Cooper SE, Oliver PL, Ponting CP (2014) The long non-coding RNA Paupar regulates the expression of both local and distal genes. Embo J 33: 296–311

Vucicevic D, Corradin O, Ntini E, Scacheri PC, Orom UA (2015) Long ncRNA expression associates with tissue-specific enhancers. Cell Cycle 14: 253–260

Wang D, Garcia-Bassets I, Benner C, Li W, Su X, Zhou Y, Qiu J, Liu W, Kaikkonen MU, Ohgi KA, Glass CK, Rosenfeld MG, Fu XD (2011a) Reprogramming transcription by distinct classes of enhancers functionally defined by eRNA. Nature 474: 390–394

Wang KC, Yang YW, Liu B, Sanyal A, Corces-Zimmerman R, Chen Y, Lajoie BR, Protacio A, Flynn RA, Gupta RA, Wysocka J, Lei M, Dekker J, Helms JA, Chang HY (2011b) A long noncoding RNA maintains active chromatin to coordinate homeotic gene expression. Nature 472: 120–124

Wang L, Park HJ, Dasari S, Wang S, Kocher JP, Li W (2013) CPAT: Coding-Potential Assessment Tool using an alignment-free logistic regression model. Nucleic Acids Res 41: e74

Welboren WJ, van Driel MA, Janssen-Megens EM, van Heeringen SJ, Sweep FC, Span PN, Stunnenberg HG (2009) ChIP-Seq of ERalpha and RNA polymerase II defines genes differentially responding to ligands. Embo J 28: 1418–1428

Werner MS, Ruthenburg AJ (2015) Nuclear Fractionation Reveals Thousands of Chromatin-Tethered Noncoding RNAs Adjacent to Active Genes. Cell Rep 12: 1089–1098

Zhang Y, Liu T, Meyer CA, Eeckhoute J, Johnson DS, Bernstein BE, Nusbaum C, Myers RM, Brown M, Li W, Liu XS (2008) Model-based analysis of ChIP-Seq (MACS). Genome Biol 9: R137

